# Neuronally sensed oxygen drives behavior and development in human-infective, skin-penetrating nematodes

**DOI:** 10.64898/2026.04.28.721222

**Authors:** Breanna Walsh, Navonil Banerjee, Gloria Bartolo, Elissa A. Hallem

**Affiliations:** Department of Microbiology, Immunology, and Molecular Genetics, University of California, Los Angeles, Los Angeles, CA 90095, USA; Molecular Biology Interdepartmental PhD Program, University of California, Los Angeles, Los Angeles, CA 90095, USA; UCLA-Caltech Medical Scientist Training Program, University of California, Los Angeles, Los Angeles, CA 90095, USA; Molecular Biology Institute, University of California, Los Angeles, Los Angeles, CA 90095, USA

## Abstract

Parasitic nematodes infect over a billion people worldwide and cause some of the most prevalent neglected tropical diseases^1-5^. Many of these parasites are skin penetrating and have both extra-host life stages that inhabit host feces and surrounding soil, and intra-host life stages that inhabit host niches such as skin, vasculature, and intestine^2,6-8^. Across life stages, these parasites encounter oxygen (O_2_) levels that range from ~21% at the soil-air interface to near-anaerobic levels in the host intestine^9-12^. However, whether parasitic nematodes detect and respond to changes in O_2_ levels was unknown. Here, we examine O_2_ sensation in skin-penetrating parasitic nematodes and find that they show robust responses to changes in O_2_ levels. Moreover, their O_2_-evoked behaviors differ from those of the free-living nematode *Caenorhabditis elegans*. We then investigate the molecular and neural mechanisms of O_2_ sensing in *Strongyloides stercoralis*, a genetically tractable human-infective nematode, and find that parasite-specific behavioral responses to O_2_ arise in part from evolutionary changes in their soluble guanylate cyclase repertoire. Finally, we find that neuronal O_2_ sensing regulates intra-host development in *S. stercoralis*. Our results demonstrate that skin-penetrating nematodes exhibit neuronally mediated O_2_ responses that are critical for multiple steps of their parasitic life cycle.

## INTRODUCTION

While soil-transmitted parasitic nematodes currently cause disease in approximately 1.5 billion people^13,14^, infection rates are expected to rise with the impacts of climate change^15-17^. This threat is particularly grave for the human-infective, skin-penetrating nematode *Strongyloides stercoralis*, which is currently estimated to infect ~610 million individuals worldwide^3^. While many infections with *S. stercoralis* are initially asymptomatic or lead to mild gastrointestinal distress, the autoinfective life cycle enables these nematodes to persist in the same human host for decades^18,19^. Due to autoinfection, strongyloidiasis can be challenging to cure with available anthelmintics (*e*.*g*., ivermectin and albendazole)^20-22^, and chronic infections can result from treatment failures^21^. Treatment efficacy is also threatened by anthelmintic resistance^23-25^, with reported cases presenting in parasitic nematodes that infect livestock^26-30^. When an untreated or undertreated host becomes immunosuppressed, unchecked propagation of *S. stercoralis* larvae can lead to hyperinfection syndrome and disseminated strongyloidiasis, of which most cases are fatal^31,32^. A better understanding of the basic biology of *S. stercoralis* is a critical prerequisite for the development of novel preventative or therapeutic strategies.

*S. stercoralis* has a complex life cycle that consists of both parasitic and free-living generations ^6,33^ (Fig. 1A). Parasitic adults reside in the host small intestine and produce young larvae, some of which develop into autoinfective larvae (aL3) that complete their life cycle in the same host, causing chronic infection, and some of which exit the host in feces. The young larvae that exit the host either develop on feces directly into infective third-stage larvae (iL3s) in the F_1_ generation, or they develop into free-living adults; the free-living adults reproduce on feces to generate an F_2_ generation comprised entirely of iL3s. The iL3 stage is a developmentally arrested larval stage homologous to the dauer stage of the free-living nematode *Caenorhabditis elegans*; iL3s must infect a host for development to continue^34,35^. iL3s actively seek hosts using host-associated sensory cues^36-43^ and, upon encountering a host, invade by penetrating through the skin^6,33^ (Fig. 1A). Throughout its life cycle, *S. stercoralis* encounters a wide range of O_2_ levels, including atmospheric levels at the soil surface, intermediate levels in host feces and vasculature, and near-anaerobic levels in the host intestinal tract^9-12,44^. This raises the possibility that O_2_ sensing may be crucial for the parasitic life cycle of *S. stercoralis*. However, while O_2_ sensing in *C. elegans* adults is well-studied^44-47^, whether *S. stercoralis* or other parasitic nematodes sense O_2_ was unknown.

**Fig. 1.**
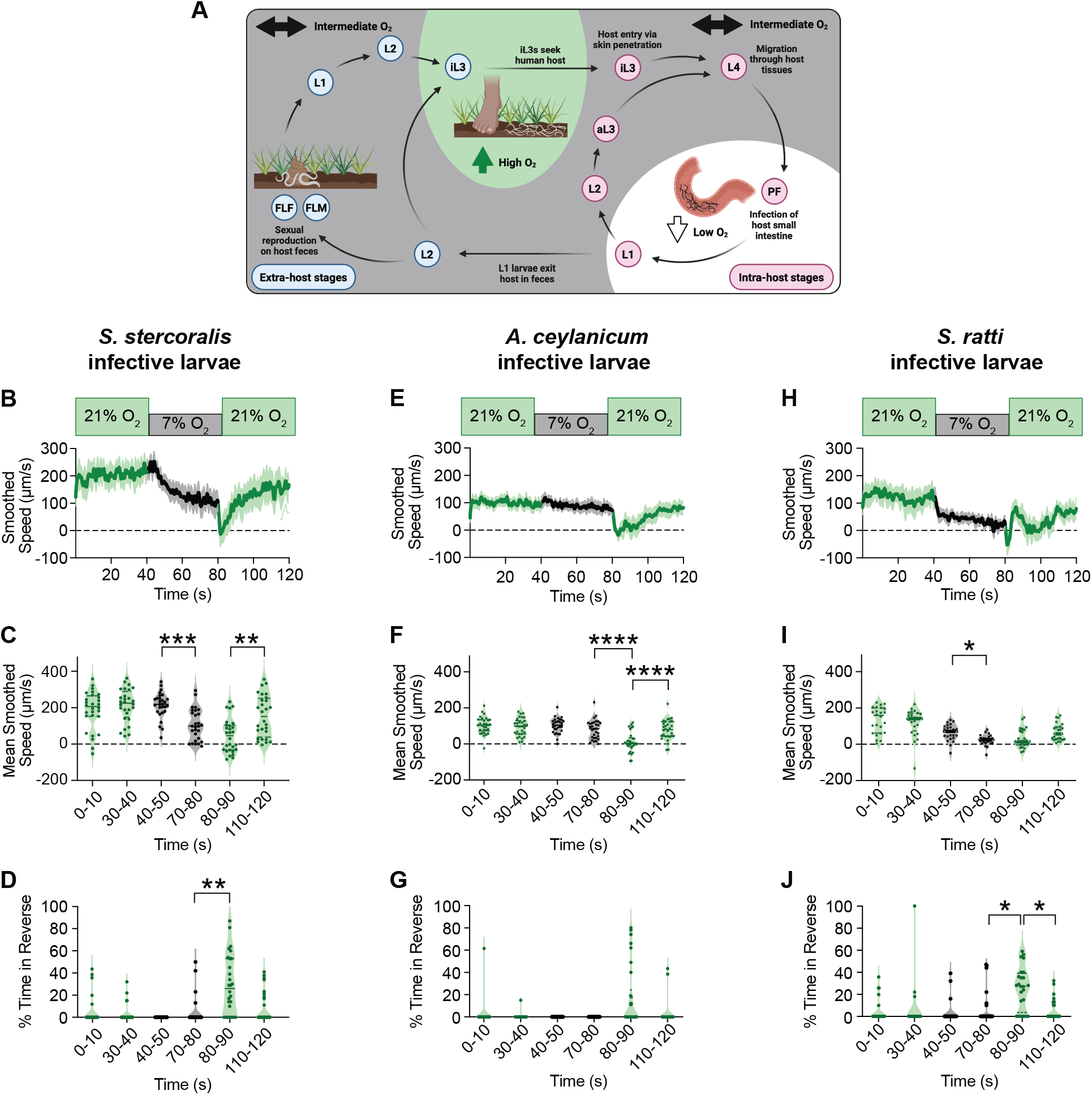
Skin-penetrating iL3s exhibit O_2_-evoked behavior. **A**. O_2_ levels vary throughout the life cycle of *S. stercoralis*^48^. Extra-host iL3s seek hosts at the soil surface^43^ (~21% O_2_^44^). iL3s enter a host via skin penetration, where they encounter intermediate, subatmospheric levels of O_2_ ^11^, and exit developmental arrest in a process called activation. They then migrate through host tissues and develop into fourth-stage larvae (L4s), which mature into parasitic females (PFs) that reside in the near-anaerobic host intestine^12,54^. Eggs laid by PFs hatch into post-parasitic first-stage larvae (L1s). Some of these L1s become autoinfective, developing into second-stage larvae (L2s) and subsequently autoinfective third-stage larvae (aL3s) that penetrate through the intestinal wall and maintain a chronic infection. Other L1s exit the host in feces and develop into post-parasitic L2s. Some of these L2s develop directly into iL3s; others develop into free-living females (FLFs) or free-living males (FLMs). FLFs and FLMs then reproduce sexually to generate post-free-living L1s. All progeny of FLFs and FLMs are obligated to develop from post-free-living L2s into iL3s, which can only continue their life cycle upon infection of a host. Non-iL3 extra-host stages inhabit feces, where O_2_ levels are subatmospheric due to respiring bacteria^40^. Pink circles depict intra-host life stages; blue circles depict extra-host life stages. O_2_ levels are noted by background color (high O_2_ in green, intermediate O_2_ in gray, low O_2_ in white). **B-D**. Response of *S. stercoralis* iL3s to acute O_2_ shifts. **B**. Graph of smoothed speed. Bold line shows mean smoothed speed; shading shows 95% confidence interval. Negative values indicate reverse movement. **C**. Violin plot of mean smoothed speed during the first and last 10 s of each gas pulse. Negative values indicate reverse movement. **D**. Violin plot showing the percentage of time spent in reverse during the first and last 10 s of each gas pulse. **E-G**. Response of *A. ceylanicum* iL3s to acute O_2_ shifts. Graphs are as described in B-D. **H-J**. Response of *S. ratti* iL3s to acute O_2_ shifts. Graphs are as described in B-D. For B-J, iL3s were exposed to a 40 s pulse of 21% O_2_ (green), followed by a 40 s pulse of 7% O_2_ (black), followed by a 40 s pulse of 21% O_2_ (green). For violin plots, dots represent individual worms, solid lines show medians, and dotted lines show quartiles. \**p*<0.05, \*\**p*<0.01, \*\*\**p*<0.001, \*\*\*\**p*<0.0001, Friedman test with Dunn’s post-test; only significant, adjacent comparisons are displayed. n = 24-30 iL3s per species.

Here, we examine O_2_ sensation in skin-penetrating parasitic nematodes. We show that *S. stercoralis* and other skin-penetrating nematodes exhibit robust motility changes when exposed to acute shifts in O_2_ concentration; these behaviors are distinct from those of *C. elegans*. We find that the genomes of *S. stercoralis* and other parasitic nematodes have a reduced repertoire of soluble guanylate cyclases (sGCs) – molecular regulators of O_2_ sensing^45^ – relative to non-parasitic nematodes such as *C. elegans*. One of these sGCs, *Sst*-GCY-35, is required for O_2_-evoked behavior and basal locomotion in *S. stercoralis* iL3s. The subset of sGCs found only in non-parasitic nematodes mediates the species-specific O_2_-evoked behaviors seen in *C. elegans*. We also identify *S. stercoralis* neurons that express *Sst*-GCY-35, detect changes in O_2_ levels, and drive O_2_-evoked behavior. Finally, we show that detection of low O_2_, mediated in part by *Sst*-GCY-35, stimulates *S. stercoralis* iL3s to exit developmental arrest, indicating a critical role for neuronally mediated O_2_ sensing in driving intra-host development in these parasites. Together, our results illuminate the neural and molecular mechanisms that enable human-parasitic worms to respond to O_2_ and highlight how evolutionary changes in the sGC repertoire enable species-specific behavioral programs. Given the robust O_2_-evoked behaviors found in parasitic nematodes, targeting the molecular machinery required for O_2_ sensing is poised as a novel strategy for infection prevention and control.

## RESULTS

### Skin-penetrating nematodes sense and respond to changes in ambient O_2_ levels

As *S. stercoralis* passes between intra-host and extra-host niches, it encounters a wide range of O_2_ concentrations^10-12,48^. Soil-surface-dwelling iL3s primarily experience atmospheric levels of O_2_ (*i*.*e*., ~21%)^11^, while parasitic females in the intestinal tract are exposed to near-anerobic conditions^12^ (Fig. 1A). In the extra-host environment, non-infective larvae and free-living adults inhabit host feces and experience intermediate O_2_levels, as aerobic fecal bacteria consume O_2_ and generate local microgradients of oxygenation^40^ (Fig. 1A). Inside a host, larval stages are similarly presented with subatmospheric levels of O_2_ that vary by tissue type (*e*.*g*., 6.5% in the end venous vasculature, 9.5% in arteries, and 13.5% in lung alveoli)^11^ (Fig. 1A). However, whether *S. stercoralis* and other skin-penetrating nematodes sense O_2_ remained a mystery.

To test if *S. stercoralis* responds to changes in ambient O_2_ levels, we quantified the motile behaviors of iL3s exposed to rapid O_2_ shifts. Using a chamber lid with a transparent viewing window^42,49,50^, we video-recorded the movement of iL3s subjected to acute O_2_ downshifts (*i*.*e*., 21% to 7% O_2_) and upshifts (*i*.*e*., 7% to 21% O_2_) and quantified motility (Fig. S1). We found that when exposed to 7% O_2_, *S. stercoralis* iL3s exhibit a gradual slowing response that is reversible when worms return to 21% O_2_ (Fig. 1B-D, S2A-C). When shifted from 7% to 21% O_2_, iL3s briefly move in reverse – a behavior tied to escape of aversive stimuli^51-53^ (Fig. 1D, S2C). These O_2_-evoked behaviors occurred regardless of whether iL3s were initially presented with an O_2_ downshift or upshift (Fig. 1B-D, S2A-C). The O_2_ downshift from 21% to 7% mimics the acute decrease in ambient O_2_ encountered by iL3s upon entering a host^11^; these data are consistent with the possibility that decreased ambient O_2_ serves as a cue marking successful host entry and allows iL3s to cease the rapid crawling necessary for host seeking.

We next asked if iL3s of other skin-penetrating nematode species could similarly respond to changes in ambient O_2_ levels. The human-infective hookworm *Ancylostoma ceylanicum* and the rat-infective nematode *Strongyloides ratti* also experience large shifts in O_2_concentration while progressing through their life cycles^11,12,48,54-57^ (Fig. S3A-B). We found that both *A. ceylanicum* and *S. ratti* iL3s were able to sense O_2_ shifts and demonstrated behaviors similar – albeit not identical – to those seen in *S. stercoralis* iL3s (Fig. 1E-J, S2D-I). While *A. ceylanicum* iL3s do not markedly change their crawling speed in the presence of 7% O_2_, they exhibit a sharp pause response immediately following the upshift to 21% O_2_ (Fig. 1E-F, S2D-E). In contrast, *S. ratti* iL3s are exceptionally responsive to shifts in O_2_, with exposure to 7% O_2_ causing most iL3s to slow to rest (Fig. 1H-I, S2G-H). Moreover, unlike *S. stercoralis* and *S. ratti* iL3s, most *A. ceylanicum* iL3s do not reverse in response to an upshift from 7% to 21% O_2_ (Fig. 1D, G, J; S2C, F, I). Together, these results demonstrate that skin-penetrating iL3s show robust behavioral responses to changes in O_2_ levels.

To determine whether O_2_-evoked behaviors in parasitic nematodes vary across life stages, we quantified the motile behaviors of *S. stercoralis* free-living females (FLFs) and free-living males (FLMs) exposed to O_2_ shifts. FLFs, like iL3s, show a decrease in speed in response to O_2_ downshifts and reverse immediately following an O_2_ upshift (Fig. S4A-F). However, this decrease occurs more rapidly in FLFs than iL3s (Fig. S4A-B, D-E). For FLFs, exposure to low O_2_ likely indicates the presence of a food source (*i*.*e*., aerobic bacteria in host feces)^40,46^ and the resultant behaviors are consistent with a preference to navigate toward and remain near food. In contrast, FLMs appear less responsive to O_2_ shifts, although they cease forward crawling immediately following an O_2_ upshift (Fig. S4G-I). For FLMs, the presence of a possible food source may be a less salient cue than pheromones amid the search for reproductively available FLFs^58^. While the acute responses of *S. stercoralis* iL3s, FLFs, and FLMs to shifts in O_2_ concentration are similar, the subtle life-stage-specific differences in these O_2_-evoked responses appear tailored for ethologically relevant behaviors.

### O_2_-evoked behaviors differ between parasitic infective larvae and *C. elegans* dauer larvae

The ability to sense O_2_ is critical for social feeding behaviors in *C. elegans* hermaphroditic adults, as it enables worms to cluster in areas where bacterial food is abundantly growing^46,59^. The O_2_-evoked motile behaviors – as well as the molecular and neuronal underpinnings of O_2_ sensing – have been well characterized in *C. elegans* adults^45-47,59-63^. However, it remains unclear if *C. elegans* dauers sense and respond to changes in ambient O_2_ concentration. As developmentally arrested third-stage larvae, the *C. elegans* dauer is the life cycle stage that is most homologous to the iL3 stage of skin-penetrating nematodes^34^.

*S. stercoralis* iL3s and *C. elegans* dauers occupy discrete ecological niches. *S. stercoralis* iL3s reside at the soil surface, engage in active host seeking, and exit developmental arrest only following host entry^43^. *C. elegans* inhabits decomposing organic matter^64^, and dauers, which develop under harsh environmental conditions (*e*.*g*., increased temperature, insufficient food), attempt to latch onto a passing insect for transport to favorable conditions^64,65^. When conditions improve – usually marked by the reintroduction of food – *C. elegans* exits the dauer stage and resumes development toward hermaphroditic adulthood^64^.

To compare the O_2_-evoked behaviors of parasitic and non-parasitic nematodes, we sought to define the O_2_-evoked motile behaviors of *C. elegans* dauers. We used the wild isolate *C. elegans* CB4856 Hawaii (HW), as this strain has the ancestral allele of *npr-1*, a gene known to regulate O_2_-mediated responses among other sensory-driven behaviors (in contrast to the laboratory N2 strain, which has a gain-of-function allele of *npr-1*)^59,66-69^. Similar to *S. stercoralis* iL3s, *C. elegans* dauers decrease their speed when exposed to 7% O_2_and increase their speed when returned to 21% O_2_ (Fig. 2A, C). Both species also reverse following an O_2_ upshift (Fig. 2B, D). However, unlike *S. stercoralis* iL3s, *C. elegans* dauers immediately pause following the downshift from 21% O_2_ to 7% O_2_ (Fig. 2A-B, E-F). This pause response is also apparent in *C. elegans* adults and absent in *S. stercoralis* FLFs (Fig. S5). The lack of an O_2_-downshift-elicited acute pause response in all tested species of parasitic nematodes (Fig. 1, S2) suggests that O_2_-evoked behavioral responses may have evolved to reinforce parasite-specific behaviors in these skin-penetrating nematodes.

**Fig. 2.**
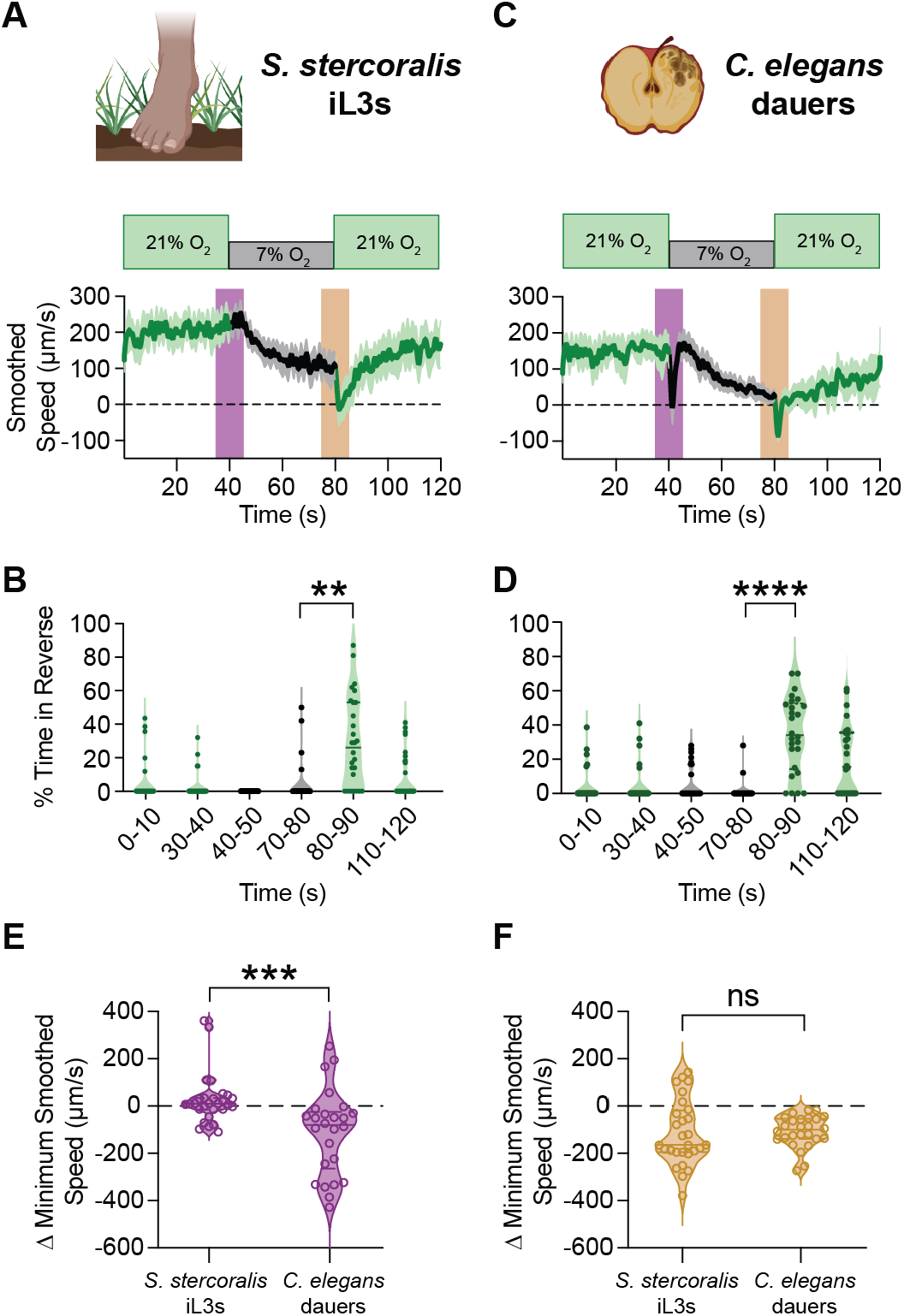
*S. stercoralis* iL3s and *C. elegans* dauers exhibit distinct behavioral responses to O_2_ shifts. **A-B**. Response of *S. stercoralis* iL3s to acute O_2_ shifts. **A**. Graph of smoothed speed. Bold line shows mean smoothed speed; shading shows 95% confidence interval. Negative values indicate reverse movement. **B**. Violin plot showing the percentage of time spent in reverse during the first and last 10 s of each gas pulse. **C-D**. Response of *C. elegans* HW dauers to acute O_2_ shifts. Graphs are as described in A-B. **E**. *C. elegans* dauers, but not *S. stercoralis* iL3s, exhibit a transient pause response following an O_2_ downshift. Violin plot shows the difference in minimum speed in the 5 s before and after the downshift (purple). **F**. *S. stercoralis* iL3s and *C. elegans* dauers respond similarly to an O_2_ upshift. For the upshift (yellow), graph is as described in E. For A-F, worms were exposed to a 40 s pulse of 21% O_2_ (green), followed by a 40 s pulse of 7% O_2_ (black), followed by a 40 s pulse of 21% O_2_ (green). For violin plots, dots represent individual worms, solid lines show medians, and dotted lines show quartiles. For B and D, \*\**p*<0.01, \*\*\*\**p*<0.0001, Friedman test with Dunn’s post-test; only significant, adjacent comparisons are displayed. For E-F, \*\*\**p*<0.001, ns = not significant, Mann-Whitney test. n = 26-28 worms per species. Data in A-B are from Fig. 1.

### The *S. stercoralis* soluble guanylate cyclase *Sst-*GCY-35 is required for O_2_-evoked behavior

We next sought to unravel the molecular and neuronal mechanisms of O_2_ sensation in parasitic nematodes. We used *S. stercoralis* for these experiments because it is unique among human-infective nematodes in that the single generation of free-living adults (Fig. 1A) provides an access point for genetic manipulation. Exogenous DNA can be introduced via intragonadal microinjection of FLFs to generate transgenics or knockouts^36,70-74^, and stable lines of transgenics or knockouts can be maintained by propagation through the laboratory host of *S. stercoralis*, the Mongolian gerbil^42,75-77^.

In *C. elegans*, soluble guanylate cyclases (sGCs) serve as molecular sensors of O_2_ within sensory neurons^46,47^. Each sGC is composed of a ligand-binding H-NOX domain, a Per-ARNT-Sim (PAS) domain, a coiled-coil domain, and a catalytic cyclase domain^46,62,78,79^. The sGCs function as dimers; O_2_ binds in the H-NOX domains of the two subunits, which catalyzes the conversion of GTP to cGMP^46,62,63,78^. In sGC-expressing sensory neurons, cGMP opens the ligand-gated cation channel formed by TAX-2 and TAX-4, resulting in cation influx^46,47^. There are seven *C. elegans* sGCs: *Cel*-GCY-31 - *Cel*-GCY-37^46,62^. Of these sGCs, *Cel*-GCY-35 and *Cel*-GCY-36 mediate motile responses to O_2_ upshifts, while *Cel*-GCY-31 and *Cel-*GCY-33 modulate behaviors tied to O_2_ downshifts^47^.

We first asked if the *S. stercoralis* genome contained homologs of the O_2_-sensing *Cel*-sGCs. We identified four candidate O_2_ sensors by protein-level homology: SSTP_0000680700, SSTP_0000411800, SSTP_0000255800, and SSTP_0000482200. Following phylogenetic analysis with *Cel-*sGCs, we named these putative *S. stercoralis* sGCs *Sst*-GCY-35, *Sst*-GCY-36, *Sst*-GCY-37.1, and *Sst*-GCY-37.2 (Fig. 3A). Using publicly available RNA-seq data^80-82^, we observed that expression levels of *Sst-gcy-35* and *Sst-gcy-36* vary with respect to life cycle stage, with high expression in extra-host iL3s and low expression in intra-host parasitic females (PFs) (Fig. 3B). In contrast, *Sst-gcy-37*.*1* expression levels appear stable with respect to life stage, while *Sst-gcy-37*.*2* shows little to no expression across life stages by RNA-seq (Fig. 3B). We focused on *Sst-gcy-35*, as this gene is expressed – albeit at disparate levels – in both iL3s and FLFs.

**Fig. 3.**
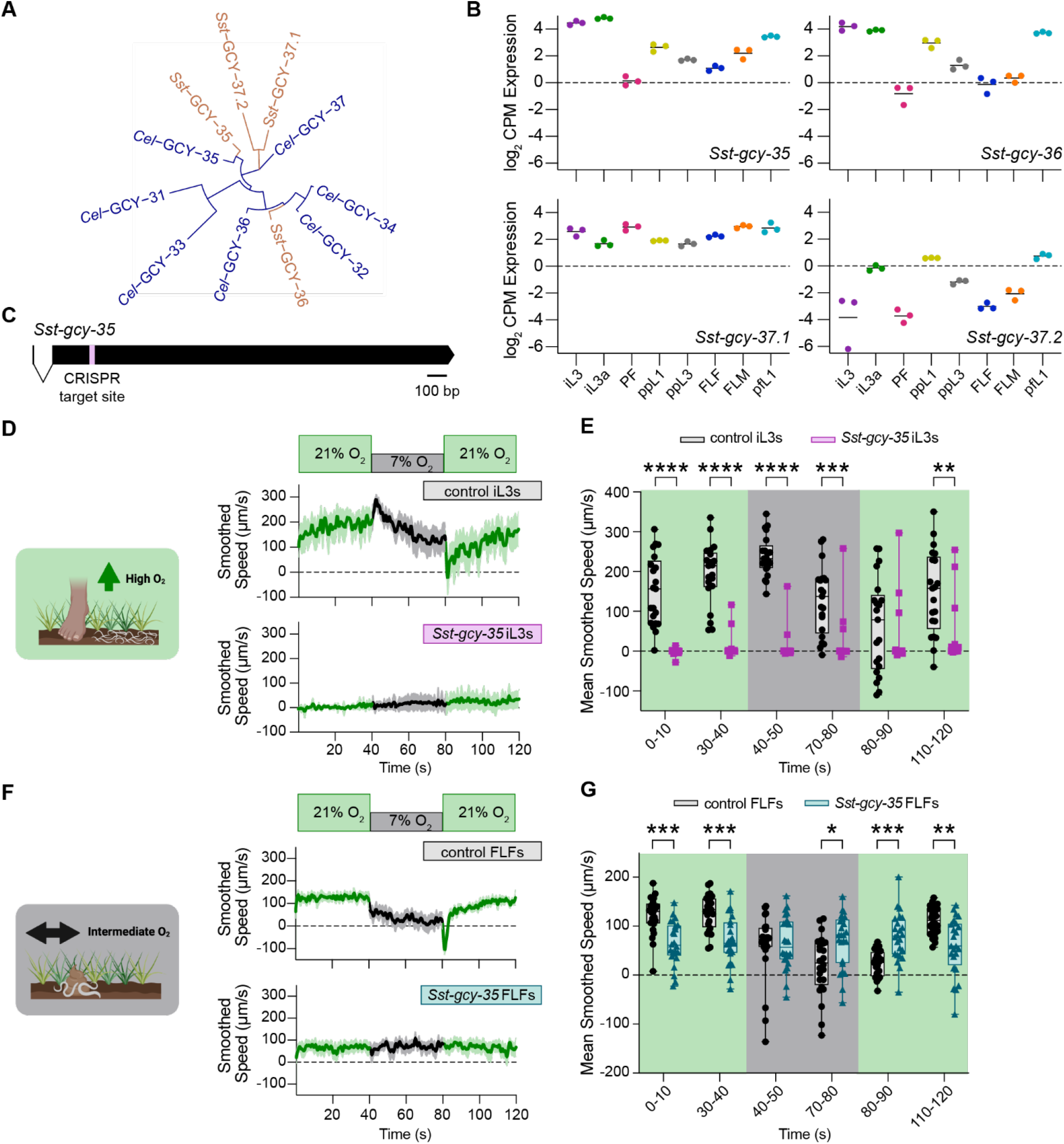
*Sst-*GCY-35 is required for O_2_-evoked behavioral responses in *S. stercoralis*. **A**. Dendrogram of *C. elegans* (blue) and *S. stercoralis* (brown) sGCs. **B**. Expression levels (in log_2_ counts per million) of the *Sst-*sGC genes across life stages based on publicly available RNA-seq data^80,81,82^. Life stages are infective larvae (iL3s), activated iL3s (iL3a), parasitic females (PFs), post-parasitic first-stage larvae (ppL1s), post-parasitic third-stage larvae (ppL3s), free-living females (FLFs), free-living males (FLMs), and post-free-living first-stage larvae (pfL1s). Each dot represents an independent replicate, with the mean shown by a solid line. **C**. Gene model of *Sst-gcy-35* showing the CRISPR target site for gene disruption. Exons are shown in black, introns are shown as black angled lines below the exons, and the CRISPR target site is shown in pink. The gene is displayed to scale; scale bar is 100 base pairs (bp). **D-E**. Response of control iL3s vs. *Sst-gcy-35* iL3s to acute O_2_ shifts. **D**. Graph of smoothed speed. Bold lines show mean smoothed speed; shading represents the 95% confidence interval. **E**. Box plot of mean smoothed speed during the first and last 10 s of each gas pulse. Each dot represents an individual worm, with the median shown by a solid line and quartiles shown by the box boundaries. Control iL3s are in gray; *Sst-gcy-35* iL3s are in pink. **F-G**. Response of control FLFs vs. *Sst-gcy-35* FLFs. Graphs are as described in D-E. Control FLFs are in gray; *Sst-gcy-35* FLFs are in blue. For D-G, worms were exposed to a 40 s pulse of 21% O_2_ (green), followed by a 40 s pulse of 7% O_2_ (black), followed by a 40 s pulse of 21% O_2_ (green). Negative values indicate reverse movement. \**p*<0.05, \*\**p*<0.01, \*\*\**p*<0.001, \*\*\*\**p*<0.0001, repeated measures two-way ANOVA with the Geisser-Greenhouse correction and Šidák’s post-test; only significant, adjacent comparisons are displayed. Note that no significant differences across time points were observed with *Sst-gcy-35* iL3s or FLFs (see Source Data File). n = 20-24 worms per combination of life stage and genotype.

We generated a stable line of *S. stercoralis* with a heritable, homozygous knockout of *Sst-gcy-35* using CRISPR/Cas9-mediated targeted mutagenesis^42,71,77,83^ (Fig. 3C, S6) and examined the behavior of *Sst-gcy-35* iL3s. We found that *Sst-gcy-35* iL3s fail to respond to either acute O_2_ upshifts or downshifts (Fig. 3D-E). In addition, *Sst-gcy-35* iL3s remain largely immobile (Fig. 3D-E); this is in stark contrast to wild-type iL3s, which crawl rapidly when exposed to 21% O_2_ (Fig. 1B-C, 3D-E). When exposed to a non-gaseous stimulus (*i*.*e*., the approximate host body temperature range of 34-37^°^C^38^), *Sst-gcy-35* iL3s exhibit rapid movement, although they remain slower than wild-type iL3s (Fig. S7). This suggests that the severe reduction in motility of *Sst-gcy-35* iL3s is not primarily due to motor deficits. Our results align with the possibility that *Sst-gcy-35* iL3s are unresponsive to 21% O_2_, and as a result exhibit a pronounced cessation in movement that mirrors the slowing seen in wild-type iL3s with prolonged exposure to low O_2_ (Fig. S2A-C). Moreover, our results suggest that the ability to sense ambient O2 at the soil surface promotes basal exploratory movement requisite for host seeking in *S. stercoralis* iL3s.

To determine whether *Sst-gcy-35* also mediates O2 responses at other life stages, we next examined the behavior of *Sst-gcy-35* FLFs. In contrast to *Sst-gcy-35* iL3s, the *Sst-gcy-35* FLFs are mobile at 21% O_2_, even at room temperature (Fig. 3D, F). When exposed to acute shifts in the O_2_ level, *Sst-gcy-35* FLFs crawl at a constant speed that is unimpacted by upshifts or downshifts in the ambient O_2_ concentration (Fig. 3F-G). The speed at which the *Sst-gcy-35* FLFs travel is slower than the speed at which wild-type FLFs crawl during exposure to 21% O_2_ (Fig. 3F-G, S4D-E). *Sst-gcy-35* FLFs also do not exhibit the upshift-associated reversals seen in wild-type FLFs (Fig. 3F-G, S4D, F). In all, these data demonstrate that *Sst-*GCY-35 is required for both *S. stercoralis* iL3s and FLFs to sense and respond to acute changes in ambient O_2_ levels.

### Differences in the sGC repertoire sculpt species-specific O_2_-evoked responses

The inability of *Sst-gcy-35* worms to respond to either downshifts and upshifts in the ambient O_2_ level was unexpected given that, in *C. elegans* adults, loss of *Cel-gcy-35* primarily impacts O_2_ upshift-associated behaviors^46,47^ (Fig. S8A-B). It was unclear, however, if similar changes in O_2_-evoked motile behaviors would appear in *C. elegans gcy-35* dauers, the life stage most homologous to parasitic iL3s^35,84^. We found that when *Cel-gcy-35* dauers are exposed to acute shifts in O_2_ concentration, the dauers – like *Sst-gcy-35* iL3s – are largely immobile at 21% O_2_ (Fig. 4A-D). These dauers remain immobile when shifted to 7% O_2_ (Fig. 4C-D). However, when *Cel-gcy-35* dauers are returned to 21% O_2_, they immediately initiate forward crawling, suggesting that *Cel-gcy-35* dauers retain the capacity to sense increases in ambient O_2_ despite lacking the known O_2_-upshift detector *Cel*-GCY-35^47^ (Fig. 4C-D). This phenotype sharply contrasts with that of *Sst-gcy-35* iL3s, which remain immobile at the O_2_ upshift (Fig. 4A-D). Thus, loss of GCY-35 results in distinct phenotypes in *S. stercoralis* and *C. elegans*.

**Fig. 4.**
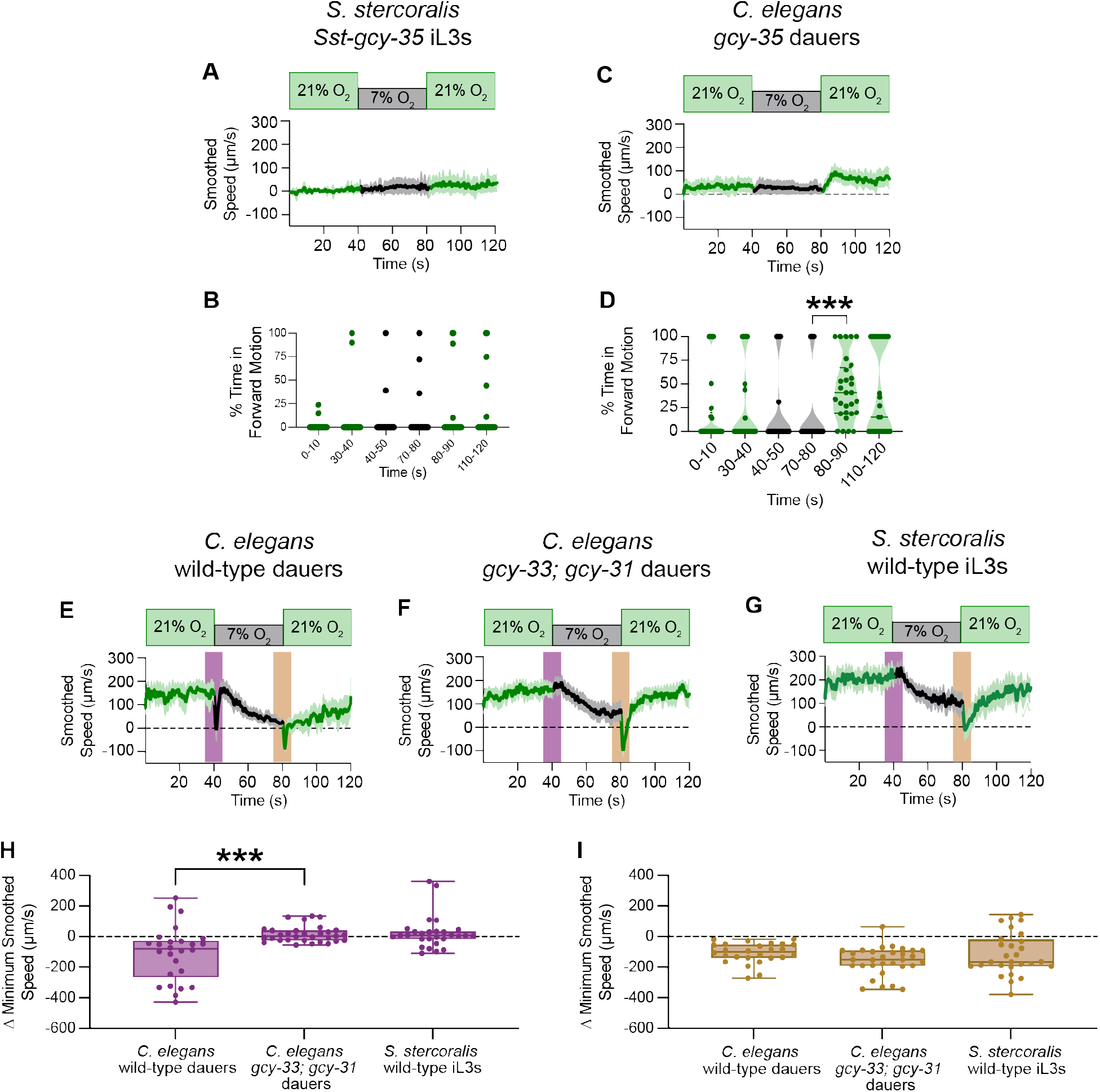
The sGC repertoire sculpts species-specific O_2_-evoked behaviors. **A-D**. *C. elegans gcy-35* dauers – but not *S. stercoralis Sst-gcy-35* iL3s - respond to O_2_ upshifts. **A, C**. Graph shows the smoothed speed. Bold lines show mean smoothed speed; shading represents the 95% confidence interval. **B, D**. Violin plot shows the percentage of time spent in forward motion during the first and last 10 s of each gas pulse. **E**. Response of wild-type *C. elegans* HW dauers to acute O_2_ shifts. Graph is as described in A, C. **F**. Response of *Cel-gcy-33; Cel-gcy-31* dauers to acute O_2_ shifts. Graph is as described in A, C. **G**. Response of *S. stercoralis* iL3s to acute O_2_ shifts. Graph is as described in A, C. **H-I**. *Cel-gcy-33; Cel-gcy-31* dauers are unable to execute O_2_-downshift-associated pause responses, while upshift-associated behaviors are preserved. Violin plots show the difference in minimum speed in the 5 s before and after the downshift (purple) or upshift (yellow). For A-I, worms were exposed to a 40 s pulse of 21% O_2_ (green), followed by a 40 s pulse of 7% O_2_ (black), followed by a 40 s pulse of 21% O_2_ (green). In A, C, and E-G, negative values indicate reverse movement. For violin plots, dots represent individual worms, with the median shown by a solid line and quartiles shown by dotted lines. For B and D, \*\*\**p*<0.001, Friedman test with Dunn’s post-test; only significant, adjacent comparisons are displayed. For H-I, \*\*\**p*<0.001, Kruskal-Wallis test with Dunn’s post-test. n = 20-31 worms per combination of species and genotype. Data in A are from Fig. 3, data in E are from Fig. 2, and data in G are from Fig. 1.

To explain these species-specific phenotypes, we hypothesized that the *C. elegans* sGCs for which there are no apparent *S. stercoralis* homologs – *Cel*-GCY-31 and *Cel*-GCY-33 – may be providing compensatory O_2_ sensation in *C. elegans* (Fig. 3A). To test this hypothesis, we first asked whether other parasitic nematodes also lacked *Cel*-GCY-31 and *Cel*-GCY-33 homologs. We searched for candidate sGCs in seventeen additional nematode species, including both parasitic and non-parasitic species in Clades I, III, IV, and V. We bioinformatically identified putative sGCs in fourteen of these species; three human-parasitic Clade I nematodes (*i*.*e*., *Ascaris lumbricoides, Trichinella spiralis*, and *Trichuris trichiura*) were, surprisingly, devoid of sGC homologs. Intriguingly, we found that most of the nematodes examined, including all of the parasitic nematodes, lack homologs of *Cel*-GCY-31 and *Cel*-GCY-33 (Fig. 5A). In *Toxocara canis* and the filarial nematodes from Clade III, only homologs to *Cel*-GCY-35 and *Cel*-GCY-36 were identified (Fig. 5A). In the free-living, insect-infective, and facultative parasitic (*i*.*e*., the life cycle is composed of free-living and mammalian-infective generations) nematodes explored in Clade IV, we exclusively found homologs to *Cel*-GCY-35, *Cel*-GCY-36, and *Cel*-GCY-37 (Fig. 5A). In Clade V, homologs to *Cel*-GCY-32, *Cel*-GCY-34, *Cel*-GCY-35, *Cel*-GCY-36, and *Cel*-GCY-37 are present in both parasitic and non-parasitic species; however, only the free-living, non-parasitic Clade V nematodes had homologs to *Cel*-GCY-31 and *Cel*-GCY-33 (Fig. 5A). Thus, the number of sGCs in these species increases with evolutionary time^85-88^ (Fig. 5B), and notably, *Cel*-GCY-31 and *Cel*-GCY-33 homologs do not appear to be present in parasitic nematodes.

**Fig. 5.**
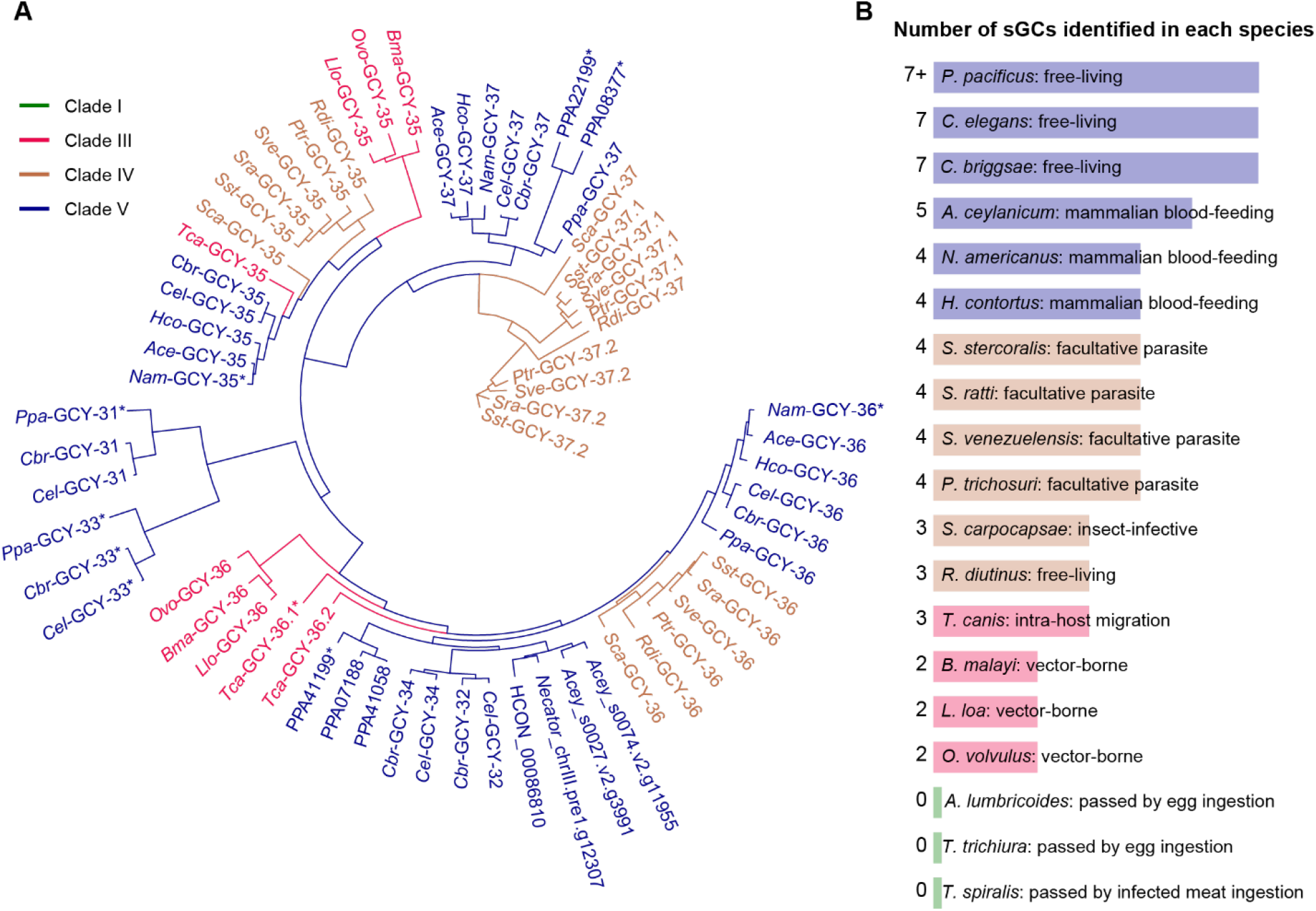
Homologs of GCY-31 and GCY-33 are limited to a subset of non-parasitic Clade V nematodes. **A**. Phylogenetic tree demonstrating relationships between candidate sGCs in representative nematodes. Clade III species (pink) include the mammalian-infective nematode *Toxocara canis* (*Tca*), the filarial nematodes *Loa loa* (*Llo*), *Brugia malayi* (*Bma*), and *Onchocerca volvulus* (*Ovo*). Clade IV species (brown) include the free-living nematode *Rhabditophanes diutinus* (*Rdi*), the entomopathogenic nematode *Steinernema carpocapsae* (*Sca*), and the skin-penetrating, facultative parasitic nematodes *S. stercoralis* (*Sst*), *S. ratti* (*Sra), Strongyloides venezuelensis* (*Sve*), and *Parastrongyloides trichosuri* (*Ptr*). Clade V nematodes (blue) include the free-living nematodes *C. elegans* (*Cel*), *Caenorhabditis briggsae* (*Cbr*), and *Pristionchus pacificus* (*Ppa*, PPA), the blood-feeding parasitic hookworms *A. ceylanicum* (*Ace*, Acey) and Necator americanus (*Nam*, Necator), and the blood-feeding, ruminant parasite *Haemonchus contortus* (*Hco*, HCON). Clade I nematodes (green) include *Ascaris lumbricoides, Trichinella spiralis*, and *Trichuris trichiura*; no sGCs were identified in these species. **B**. The number of identified sGCs in each species increases with evolutionary time. The size of the colored bar for each species corresponds to the number of identified sGCs. Bar color corresponds to nematode clade (I – green, III – pink, IV – brown, V – blue).

In *C. elegans*, sGCs act in distinct sensory neurons to detect changes in ambient O_2_ levels. Specifically, *Cel*-GCY-35 and *Cel*-GCY-36 act in URX, AQR, and PQR neurons to detect stepwise increases in O_2_, while *Cel*-GCY-31 and *Cel*-GCY-33 act in BAG neurons to detect stepwise decreases in O_2_ ^47,51,63,89^. To disentangle the roles of these sGCs in *C. elegans* dauers, we exposed dauers lacking *Cel-gcy-33* and *Cel-gcy-31*, as well as dauers lacking *Cel-gcy-35* and *Cel-gcy-36*, to acute O_2_ shifts. Similar to wild-type dauers, dauers lacking *Cel-gcy-33* and *Cel-gcy-31* gradually slow with exposure to 7% O_2_ and reverse upon encountering an upshift in the O_2_ concentration (Fig. 4E-F). However, in contrast to wild-type dauers, they fail to show a downshift-associated acute pause response (Fig. 4E-F, 4H). Remarkably, without *Cel*-GCY-31 and *Cel*-GCY-33, *C. elegans* dauers exhibit O_2_-mediated motile responses that mimic those of wild-type *S. stercoralis* iL3s (Fig. 4F-I). Dauers lacking *Cel-gcy-35* and *Cel-gcy-36* phenocopy dauers lacking only *Cel-gcy-35*, consistent with the theory that *Cel*-GCY-35 and *Cel*-GCY-36 bind O_2_ as a heterodimer^63^ (Fig. S8C-F). Together, these results strongly suggest that species-specific O_2_-evoked behaviors arise from evolutionary changes in the sGC repertoire, and that *S. stercoralis* and other parasitic nematodes do not pause following an O_2_downshift because they lack *Cel*-GCY-31 and *Cel*-GCY-33 homologs.

### *S. stercoralis* sGCs are expressed in putative O_2_-sensing neurons that contribute to O_2_-evoked behaviors

To establish the neuronal basis of O_2_ sensation in *S. stercoralis*, we first sought to identify the O_2_-sensing neurons in these worms. Using transcriptional reporters for the *S. stercoralis* sGCs, we found that *Sst-gcy-35, Sst-gcy-36*, and *Sst-gcy-37*.*1* are expressed in a subset of head and tail neurons that bear anatomic and positional similarity to the *C. elegans* URX, AQR, and PQR neurons^46,47,60,61,90^ (Fig. 6A, S9). We were unable to detect expression from the transcriptional reporter for *Sst-gcy-37*.*2* in iL3s, which aligns with the negligible expression levels across life stages reported by RNA-seq (Fig. 3B). Based on position and molecular identity, we named the sGC-expressing neurons of *S. stercoralis* the *Sst*-URX, *Sst*-AQR, and *Sst*-PQR neurons. To test the role of these neurons in mediating O_2_-evoked behaviors, we silenced *Sst*-URX in *S. stercoralis* iL3s and exposed these worms to shifts in the ambient O_2_ concentration; we focused on the URX neurons because of their important role in driving O_2_-evoked behavior in *C. elegans*^46,47^. To achieve neuronal silencing of *Sst*-URX, we expressed the light chain of tetanus toxin (TeTx)^68^ specifically under the control of the *Sst-gcy-36* promoter; when expressed, TeTx blocks vesicle fusion at the synapse and precludes neurotransmitter release^42,68,91^. *Sst*-URX-silenced iL3s exhibit dampened responses to changes in ambient O_2_ (Fig. 6B-C), suggesting that *Sst*-URX contributes to O_2_-evoked motile behaviors.

**Fig. 6.**
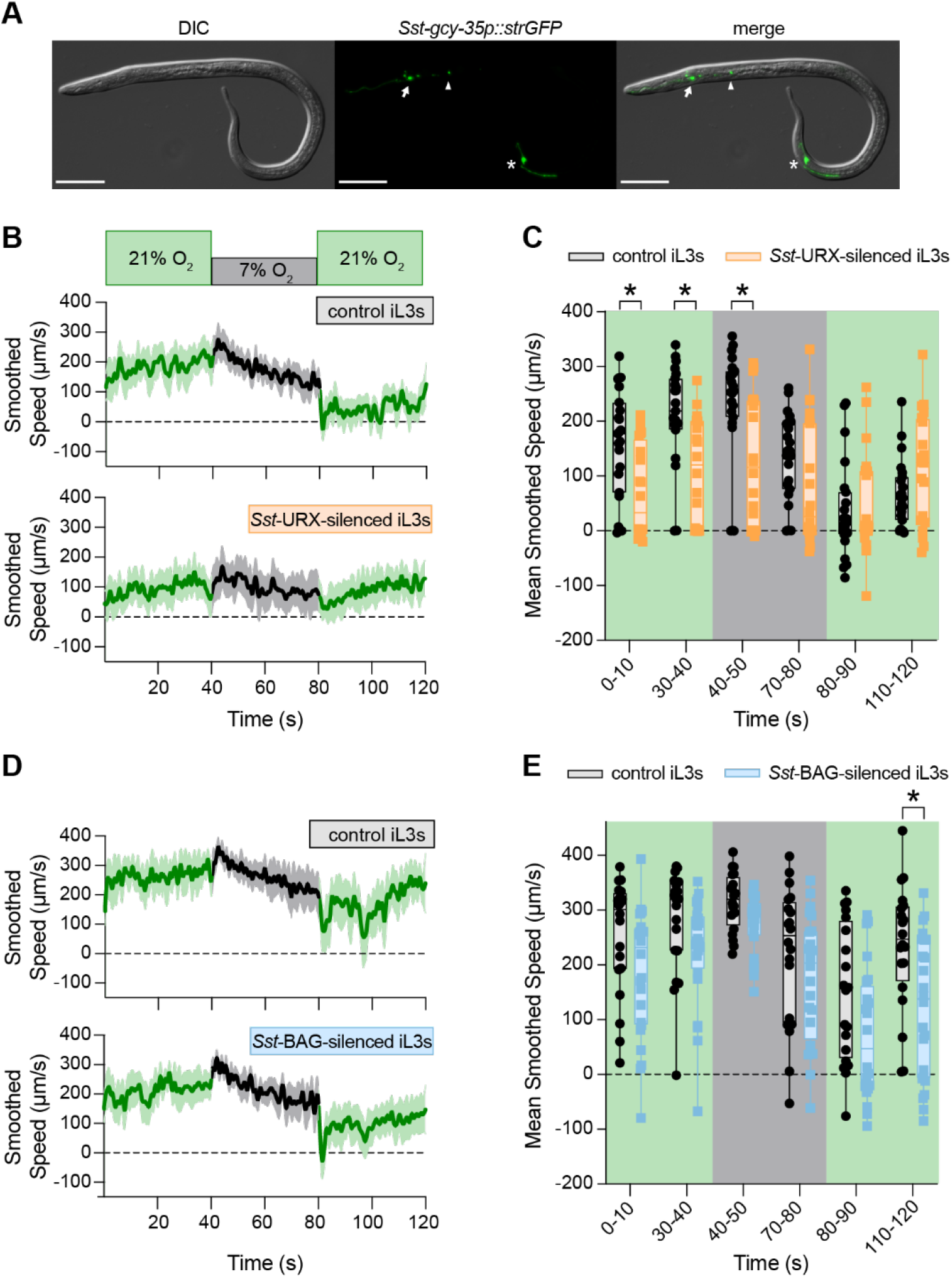
*Sst-*URX neurons, but not *Sst*-BAG neurons, contribute to O_2_-evoked behaviors. **A**. *Sst-gcy-35* is expressed in a subset of head and tail neurons that are anatomically and positionally similar to the O_2_-sensing neurons in *C. elegans*. Left panel, differential interference contrast (DIC) image; middle panel, GFP expression driven by the *Sst-gcy-35* promoter (green); right panel, merge. Arrow, arrowhead, and asterisk indicate the URX, AQR, and PQR neurons, respectively. Head is to the left; scale bar is 50 µm. **B-C**. Silencing *Sst*-URX neurons dampens O_2_-evoked behavior. **B**. Graphs show smoothed speeds of control (top, gray) vs. *Sst*-URX-silenced (bottom, orange) iL3s to acute O_2_ shifts. Bold lines show mean smoothed speed; shading represents the 95% confidence interval. **C**. Box plot shows mean smoothed speeds of control (gray) vs. *Sst*-URX-silenced (orange) iL3s during the first and last 10 s of each gas pulse. Dots represent individual worms, with the median shown by a solid line and quartiles shown by box boundaries. **D-E**. Silencing *Sst*-BAG neurons does not dampen O_2_-evoked behavior. **D**. Graphs show control (top, gray) vs. *Sst*-BAG-silenced (bottom, blue) to acute O_2_ shifts. Graph is as described in B. **E**. Box plot shows smoothed speeds of control (gray) vs. *Sst*-BAG-silenced (blue) iL3s during the first and last 10 s of each gas pulse. Graph is as described in C. For B-E, iL3s were exposed to a 40 s pulse of 21% O_2_ (green), followed by a 40 s pulse of 7% O_2_ (black), followed by a 40 s pulse of 21% O_2_ (green). Negative values indicate reverse movement. \**p*<0.05, repeated measures two-way ANOVA with the Geisser-Greenhouse correction and Šidák’s post-test; only significant, adjacent comparisons are displayed. n = 17-23 worms per genotype and condition.

In *C. elegans*, the BAG neurons sense both carbon dioxide (CO_2_) and O_2_, and mediate CO_2_ responses as well as responses to O_2_ upshifts^47,51,92-94^. We previously identified the *Sst*-BAG neurons as the primary CO_2_-sensing neurons in *S. stercoralis*^42^, but whether the *Sst*-BAG neurons also respond to O_2_ was unknown. We found that when *Sst*-BAG is silenced using TeTx in *S. stercoralis* iL3s, the worms exhibit O_2_-evoked responses similar to those seen in wild-type iL3s (Fig. 6D-E). Thus, while *C. elegans* BAG neurons mediate responses to both CO_2_ and O_2_, *S. stercoralis* BAG neurons appear to mediate responses to CO_2_ but not O_2_. Together, these results suggest that O_2_-evoked responses in free-living and parasitic nematodes are mediated by distinct but overlapping sets of sensory neurons.

### The *Sst-*URX neurons, but not the *Sst*-BAG neurons, encode responses to O_2_shifts

To determine how changes in ambient O_2_ level are neuronally encoded in *S. stercoralis*, we monitored the activity of the *Sst*-URX neurons using *in vivo* calcium imaging. In *C. elegans, Cel*-URX is inhibited by O_2_ downshifts and activated by O_2_ upshifts^47^. To measure O_2_-evoked responses in *Sst*-URX, we used a polycistronic vector to simultaneously express *Strongyloides*-codon-optimized versions of the genetically encoded calcium indicator GCaMP7f (*strGCaMP7f*)^95^ and mScarlet-I (*strmScarlet-I*) under control of the *Sst-gcy-36* promoter (Fig. 7A). We then recorded fluorescent signals from GCaMP7f and mScarlet-I while exposing iL3s to acute shifts in O_2_ concentration; changes in GCaMP7f fluorescence intensity were normalized to mScarlet-I fluorescence to capture ratiometric measurements of neuronal activity. When transitioned from 21% to 7% O_2_, *Sst*-URX neurons are inhibited (Fig. 7B-D). This inhibition is sustained for the entire duration (40 s) of 7% O_2_ exposure, suggesting that low levels of atmospheric O_2_ might cause tonic inhibition of *Sst*-URX. At the upshift in O_2_ concentration from 7% to 21%, *Sst*-URX neurons display a sharp increase in neuronal activity that slowly decays (Fig. 7B-C, E). We performed air control experiments to rule out the impact of mechanical stimulation on neuronal responses; *Sst*-URX responses to successive pulses of 21% O_2_ result in neuronal activity that is unchanged from baseline (Fig. S10A-B). These data show that *Sst*-URX is a chemosensory neuron pair that detects changes in the O_2_concentration in *S. stercoralis*.

**Fig. 7.**
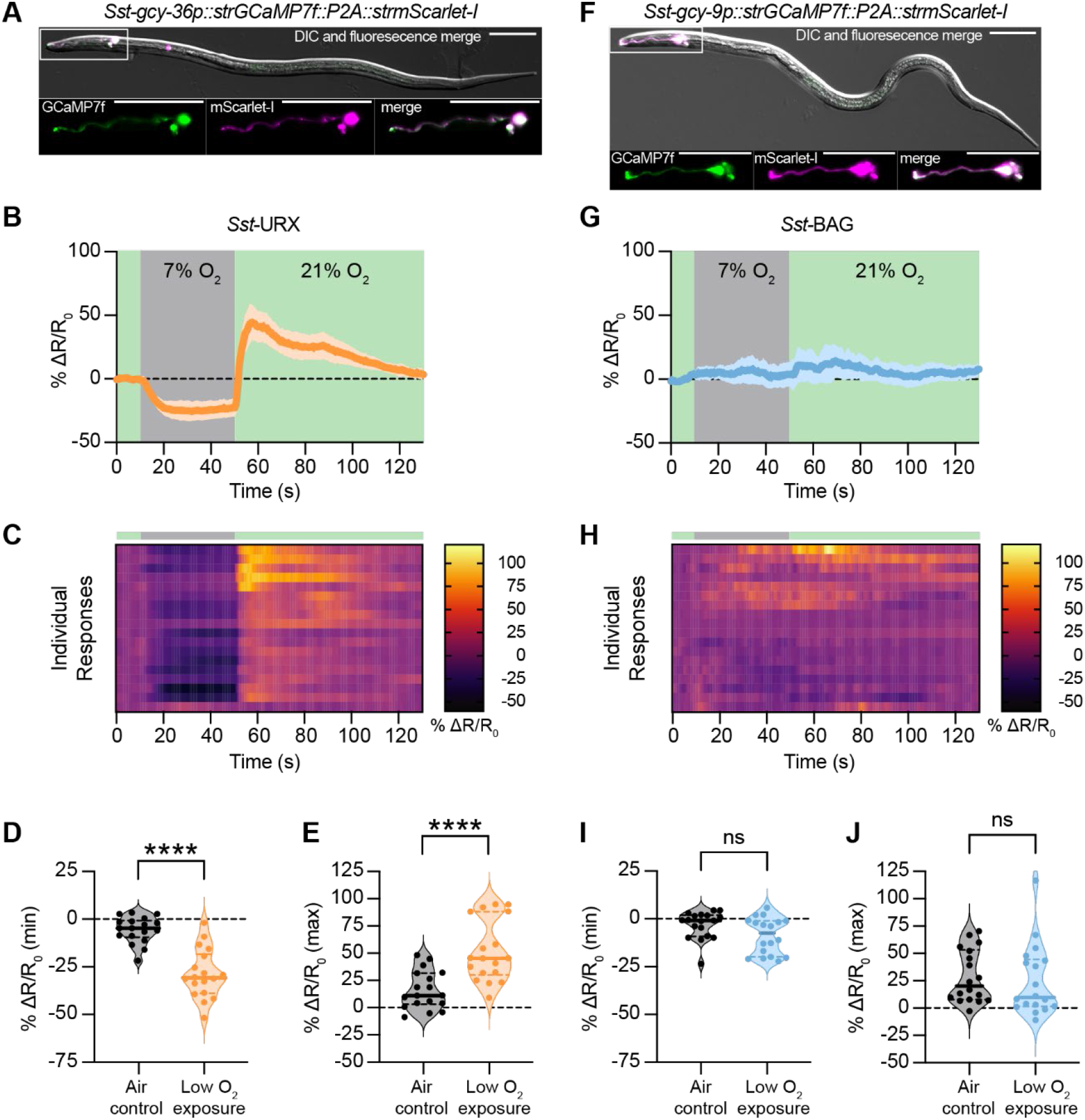
*Sst-*URX neurons, but not Sst-BAG neurons, encode responses to O_2_ shifts. **A**. An iL3 expressing an *Sst-gcy-36p::strGCaMP7f::P2A::strmScarlet-I* transgene, in which *Strongyloides*-codon-optimized (*str*) genes encoding GCaMP7f and mScarlet-I are expressed in *Sst-*URX. Top image shows DIC and fluorescence merge; lower images show GCaMP7f (left), mScarlet-I (middle), and merge (right). Head is to the left; scale bar = 50 µm. **B-E**. The *Sst-*URX neurons exhibit inhibition when exposed to 7% O_2_, followed by an increase in activity when shifted to 21% O_2_. Worms were exposed to a 40 s pulse of 21% O_2_ (green), followed by a 40 s pulse of 7% O_2_ (gray), followed by an 80 s pulse of 21% O_2_ (green); only the last 10 s of the initial pulse are displayed (t = 0-10 s). **B**. Graph shows the change in fluorescence intensity relative to baseline (% ΔR/R_0_); bold lines show mean responses, and shading represents the 95% confidence interval. **C**. Heatmap shows the neuronal responses of individual worms (rows). Bar indicates the stimulus paradigm. **D**. The minimum responses of *Sst-*URX from t = 10-50 s (orange) are decreased in worms exposed to 7% O_2_, when compared to minimum responses from control worms exposed to 21% O_2_ (black). **E**. The maximum responses of *Sst-*URX from t = 50-90 s (orange) are increased in worms exposed to 7% O_2_, when compared to maximum responses from control worms exposed to 21% O_2_ (black). \*\*\*\**p*<0.0001, unpaired t test with Welch’s correction. For D-E, dots represent individual worms, solid lines indicate medians, and dotted lines indicate interquartile ranges. **F**. An iL3s expressing an *Sst-gcy-9p::strGCaMP7f::P2A::strmScarlet-I* transgene, in which *Strongyloides*-codon-optimized genes encoding GCaMP7f and mScarlet-I are expressed in *Sst-*BAG. Image is as described in A. **G-J**. The *Sst-*BAG neurons do not exhibit O_2_-evoked activity. Stimulus paradigm and graphs are as described in B-E. ns = not significant, Mann-Whitney test. n = 18 worms per genotype.

We also imaged from the *S. stercoralis* BAG neurons of iL3s in transgenic animals that expressed *strGCaMP7f* and *strmScarlet-I* in *Sst*-BAG under the control of the *Sst*-BAG-specific *Sst-gcy-9* promoter (Fig. 7F). Unlike the *Sst*-URX neurons, the *Sst*-BAG neurons did not respond to changes in the ambient O_2_ concentration (Fig. 7G-J), with responses mimicking those seen in air controls (Fig. S10C-D). While the *Sst*-BAG neurons failed to respond to changing O_2_ levels, exposure to high concentrations of CO_2_ induced strong, rapid responses in *Sst*-BAG (Fig. S10E-H), validating the efficacy of the calcium indicator in these neurons. Thus, unlike in *C. elegans*, the *Sst-*BAG neurons do not appear to be involved in O_2_ sensation in *S. stercoralis*, likely because they lack *Sst-*sGC O_2_ sensors that are homologous to those found in *Cel*-BAG. These results highlight key differences in chemosensory neuron function across nematode species, despite the general conservation of nematode sensory neuroanatomy^96-104^.

### O_2_ sensing regulates developmental activation in *S. stercoralis*

When iL3s invade a host, they experience a sudden drop in O_2_ levels from ~21% O_2_ at the soil surface to ~7% in host tissue^11,44^. Whether detection of the acute drop in O_2_ upon host entry is one of the sensory cues that drives exit from the developmentally arrested iL3 stage, a process called activation^105,106^, was unknown. To test this possibility, we performed *in vitro* activation assays in which we exposed either wild-type or *Sst-gcy-35* iL3s to host-like conditions (DMEM media, 37°C, and 5% CO_2_)^41,42,105,106^ at either 7% O_2_ or 21% O_2_ and then measured activation frequencies (Fig. 8A). We found that wild-type iL3s display an increase in activation frequency at 7% O_2_ compared to 21% O_2_ (Fig. 8B). The *Sst-gcy-35* iL3s also show increased activation at 7% O_2_, although this increase is less pronounced than the increase observed with wild-type iL3s (Fig. 8B). Thus, low O_2_ levels stimulate *S. stercoralis* iL3s to exit developmental arrest. In addition, we found that *Sst-gcy-35* iL3s activate at a higher frequency than wild-type iL3s at 21% O_2_ but not 7% O_2_ (Fig. 8B). These results suggest that low-O_2_-evoked activation is at least partially mediated by neuronal O_2_ detection. These data align with the possibility that – independent of the ambient O_2_ concentration – the defect in O_2_ sensing in *Sst-gcy-35* iL3s mimics a low O_2_ state, thus promoting higher activation rates than those seen in wild-type iL3s. Thus, detection of low O_2_ upon host entry appears to play an important and previously unrecognized role in driving intra-host development. Interestingly, the finding that not only wild-type iL3s but also *Sst-gcy-35* iL3s activate at a higher frequency at 7% O_2_ than 21% O_2_ suggests that either neuronal detection of low O_2_ by *Sst-gcy-35*-independent mechanisms (*e*.*g*., by other *Sst-*sGCs) and/or physiological effects of low O_2_ also contribute to iL3 activation.

**Fig. 8.**
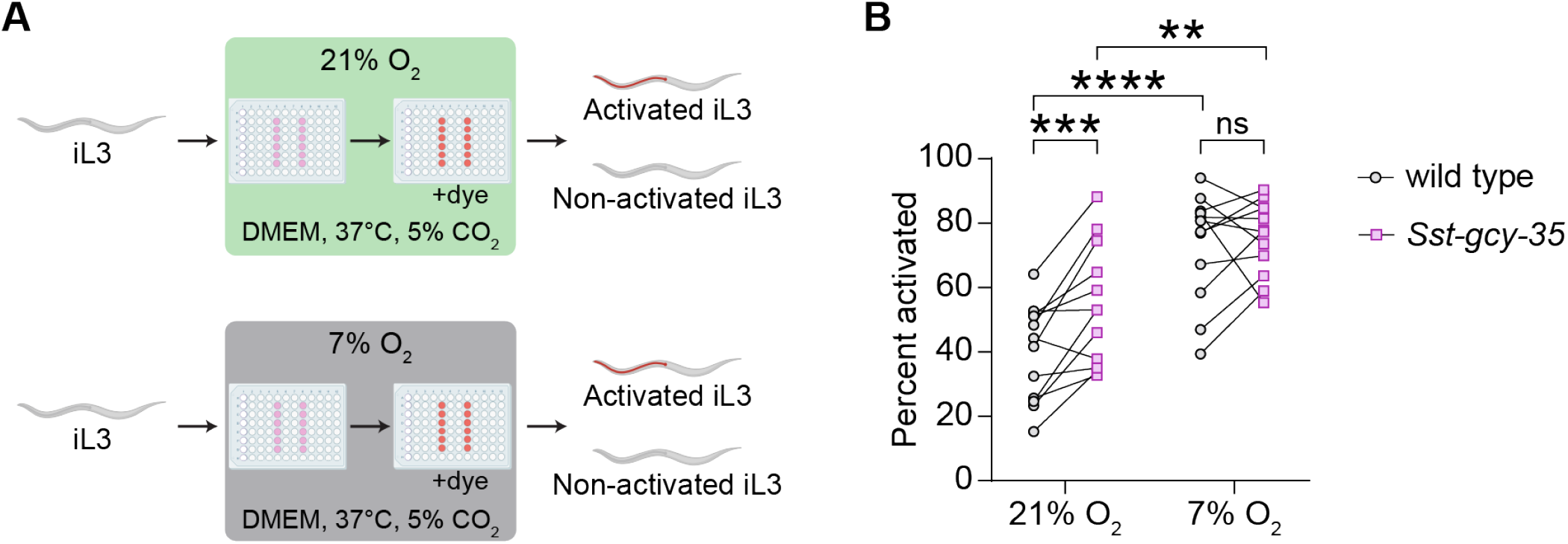
Detection of low O_2_ levels stimulates activation. **A**. Schematic of the *in vitro* activation assay. iL3s were exposed to host-like conditions (DMEM, 37°C, 5% CO_2_) at either 21% O_2_ (green) or 7% O_2_ (gray). To determine activation frequencies, the iL3s were then incubated with red fluorescent dye, and activated iL3s were identified based on the presence of red fluorescence in their pharynxes, which indicates the resumption of feeding that occurs upon activation^42^. **B**. For both wild-type and *Sst-gcy-35* iL3s, activation frequencies were lower at 21% O_2_ than 7% O_2_. In addition, *Sst-gcy-35* iL3s activated at a higher frequency than wild-type iL3s at 21% O_2_ but not 7% O_2_. \*\**p*<0.01, \*\*\**p*<0.001, \*\*\*\**p*<0.0001, ns = not significant, two-way repeated measures ANOVA with Fisher’s LSD test pairing assays performed concurrently, since these assays were statistically correlated (*r* = 0.7295, *p*<0.01, Pearson’s test). n = 12 trials per condition, with at least 200 iL3s in each condition in each trial.

## DISCUSSION

Here, we show that the human-parasitic, skin-penetrating nematode *S. stercoralis* detects and responds to changes in ambient O_2_ levels. Moreover, neuronally sensed O_2_ regulates behavior at multiple steps of the parasitic life cycle, including environmental navigation by iL3s and free-living adults as well as exit from developmental arrest by iL3s. Both behavioral and developmental responses of *S. stercoralis* to O_2_ require *Sst-*GCY-35. Notably, *Sst-gcy-35* iL3s remain nearly immobile in the absence of an applied stimulus, suggesting that O_2_sensing is necessary for the exploratory locomotion that characterizes host seeking. Our results demonstrate a previously unrecognized role for O_2_sensing in shaping the multifaceted interactions of human-parasitic nematodes with their hosts.

The skin-penetrating nematodes *S. ratti* and *A. ceylanicum* also show O_2_-evoked motile behaviors akin to those of *S. stercoralis*. The hookworm *A. ceylanicum* is distantly related to *S. stercoralis* and *S. ratti*; hookworms and *Strongyloides* species occupy distinct clades, and phylogenetic analysis indicates that the ability to invade hosts via skin penetration evolved independently as a result of convergent evolution^86,87^. Thus, their similar O_2_-evoked locomotory behaviors suggest an important role for these behaviors in successful parasitism. The increased motility of hookworm and *Strongyloides* iL3s at 21% O_2_ may reflect the need for rapid navigation toward hosts at the soil surface, whereas the slowing response to a drop in O_2_ levels may reflect the need to reduce crawling speed following entry into the low O_2_ environment of host tissue^10,11,48^.

A comparison of the O_2_-evoked behaviors of skin-penetrating nematodes with those of *C. elegans* provides insight into the evolutionary adaptations that generate species-specific sensory behaviors. Nematodes are the most abundant animals on Earth and inhabit environments ranging from the Siberian permafrost to the human intestine^2,8,48,87,107-109^. Remarkably, many of these nematode species share conserved sensory neuroanatomy^96-104^, raising the question of how species-specific sensory adaptations that enable survival in such diverse environments can arise within the constraints of an anatomically conserved sensory nervous system. We have found that, in the case of O_2_ sensing, species-specific adaptations arise from evolutionary changes in the size of the sGC repertoire. Indeed, our analysis demonstrates that the number of sGCs present in different nematodes increases with evolutionary time. Concordantly, species appear to expand their sGC repertoire in alignment with the number of distinct habitats they encounter throughout their life cycle. This suggests that robust O_2_ sensing is requisite for successful navigation of heterogeneous environments.

We also found that *S. stercoralis* lacks homologs of the *C. elegans* O_2_-downshift detectors *Cel*-GCY-31 and *Cel*-GCY-33, and as a result lacks the acute pause response to O_2_ downshifts exhibited by *C. elegans*. Moreover, phylogenetic analysis revealed that, across 19 species of nematodes, only free-living clade V nematodes possess homologs to the *C. elegans* O_2_-downshift detectors. Skin-penetrating parasitic nematodes likely lack these receptors because an acute pause response to low O_2_ would prevent the forward movement necessary for entry into the hypoxic environment of host skin. More generally, our results suggest that the tuning of the specific sGCs in a nematode’s repertoire may allow for species-specific behaviors. For example, it is possible that expansion of GCY-32 and GCY-34 homologs in *H. contortus, A. ceylanicum*, and *N. americanus* may support blood-feeding behaviors, whereas GCY-37 homologs in facultative parasites could allow for switching between free-living and parasitic generations. Future characterization of the full sGC repertoire of *S. stercoralis* and other parasitic nematodes could address these possibilities.

At the neuronal level, we find that evolutionary changes in the sGC repertoire result in differential functional engagement of anatomically conserved sensory neurons. Previous studies of anatomically conserved sensory neurons across species have revealed that many share a common sensory modality – for example, the ASE neurons mediate salt chemotaxis in both free-living and parasitic nematodes, while the AFD neurons mediate temperature sensing^39,110-113^. In the case of O_2_ sensing, the URX neurons detect O_2_ in both *C. elegans*^47^ and *S. stercoralis*. However, our results suggest that the BAG neurons detect both O_2_ and CO_2_ in *C. elegans*^47^ but only CO_2_ in *S. stercoralis*, presumably due to the lack of *Cel*-GCY-31 and *Cel*-GCY-33 homologs. Thus, sensory modalities are not universally conserved across homologous nematode sensory neurons but can be species-specific. The extent to which species-specific sensory behaviors also arise from changes in downstream circuity remains to be determined.

In addition to stimulating O_2_-evoked behaviors in skin-penetrating nematodes, shifts in ambient O_2_ levels serve as salient developmental cues. We find that activation of iL3s is enhanced at low ambient O_2_, in line with the low O_2_ environment experienced by iL3s inside the host. Moreover, *Sst*-GCY-35 contributes to the O_2_-dependent regulation of activation, such that *Sst-gcy-35* iL3s activate at a higher frequency than wild-type iL3s. Together, these results suggest that neuronally mediated detection of the O_2_ downshift that occurs upon host entry is an important cue for intra-host development. Moreover, these results have implications for a known gap in the current treatment regimens for strongyloidiasis: while ivermectin and albendazole are effective at clearing adult worms, autoinfective larvae are sometimes spared, leading to treatment failure^20,114-116^. Given that the sensation of low O_2_ promotes activation, it is possible that drugs that inhibit nematode sGCs could be used with ivermectin as part of a “shock and kill” strategy similar to that proposed for latent HIV infections^117,118^ – such drugs could cause autoinfective larvae to activate, thereby increasing their susceptibility to ivermectin.

Together, our findings demonstrate that skin-penetrating nematodes use O_2_ as a vital sensory cue. Moreover, neuronal O_2_ sensing facilitates both parasite-specific motile behaviors and intra-host development, suggesting a critical role for the O_2_-sensing pathway in enabling successful infection of a human host. Interruption of O_2_ sensing is now poised as a promising tactic for helminth control.

## METHODS

### Ethics statement

All animal protocols and procedures were approved by the UCLA Office of Animal Research Oversight (Protocol ARC-2011-060). The protocol follows the guidelines set by the AAALAC and the *Guide for the Care and Use of Laboratory Animals*.

### Strains

The following strains were used in this study: *S. stercoralis* UPD, *S. stercoralis* EAH467 *Sst-gcy-35*(*bru7*[*Sst-act-2p::strGFP::Sst-era-1* 3’ UTR]) / *Sst-gcy-35*(*bru8*[*Sst-act-2p::strmScarlet-I::Sst-era-1* 3’ UTR]) II, *A. ceylanicum* Indian strain (US National Parasite Collection Number 102954), *S. ratti* ED231, *C. elegans* CB4856 Hawaii (HW), *C. elegans* PHX9398 *gcy-33(syb9398)* V; *gcy-31(syb9309)* X, *C. elegans* PHX9450 *gcy-35a(syb9451)* I, and *C. elegans* PHX9575 *gcy-35a(syb9451)* I; *gcy-36(syb9574)* X. *C. elegans* strains PHX9450, PHX9398, and PHX9575 were generated by SunyBiotech (Fuzhou, Fujian, China) using *C. elegans* CB4856 HW as the background strain.

### Maintenance of the *Strongyloides stercoralis* UPD strain

Mongolian gerbils (Charles River Laboratories) served as the laboratory hosts for serially passaged *S. stercoralis*, as previously described^72^. In brief, infections were established by subcutaneous injection of 8-12 gerbils with an infective dose of ~2000 iL3s/gerbil in ~200 µL of sterile phosphate buffered saline (PBS). During the patency period (*i*.*e*., 14-44 days post-injection), infected gerbils were placed overnight on wire-mesh racks in cages lined with damp cardboard (Shepherd Techboard, 8 × 16.5 inches, Newco, 999589). Each morning, fecal pellets deposited overnight on the cardboard were collected, mixed with autoclaved charcoal (Bone char, 10 × 28 mesh, Ebonex) and dH_2_O, and packed into 10 cm Petri dishes (ThermoFisher, 07-000-684) lined with damp Whatman paper. Secured in plastic bins lined with damp paper towels, fecal-charcoal plates were kept at 20°C for two days and then transferred to 23°C for long-term storage. Free-living adults were isolated from fecal-charcoal plates kept either at 20°C for two days or 25°C for one day, using a Baermann apparatus^72^. iL3s were isolated from 7-10 day old fecal-charcoal plates using a Baermann apparatus^72^.

### Maintenance of the *Strongyloides stercoralis* EAH467 strain

*S. stercoralis* EAH467 was maintained in Mongolian gerbils (Charles River Laboratories). *S. stercoralis* EAH467 iL3s – either freshly isolated from a fecal-charcoal plate or following *in vitro* activation^119^ – were introduced to gerbils via oral gavage^42,75^. *S. stercoralis* EAH467 iL3s were activated using host-like conditions (*i*.*e*., incubation in Dulbecco’s Modified Eagle’s Medium [Gibco, 11995-065], at 37°C with 5% CO_2_) as previously described^42,75,119^ for 40-48 h. ~100-500 iL3s or activated iL3s were resuspended in 200 µL of sterile PBS and inoculated into each gerbil. Fecal collections and the generation of coprocultures were performed as described for the *S. stercoralis* UPD strain. *S. stercoralis* EAH467 FLFs were isolated from fecal-charcoal plates incubated at 25°C for one day; *S. stercoralis* EAH467 iL3s were isolated from fecal-charcoal plates incubated at 23°C for 7-10 days.

### Maintenance of the *Strongyloides ratti* ED321 strain

*S. ratti* was maintained in Sprague-Dawley rats (Inotiv), as previously described^72^. Infections were established by subcutaneous injection of 2-4 rats with an infective dose of ~700 iL3s/rat in ~200 µL of sterile PBS. During the patency period (*i*.*e*., 7-21 days post-injection), fecal pellets were collected and mixed into coprocultures as described for the *S. stercoralis* UPD strain. Fecal-charcoal plates were kept at 20°C for two days and then transferred to 23°C for long-term storage. iL3s were isolated from 7-10-day-old fecal-charcoal plates using a Baermann apparatus^72^.

### Maintenance of *Ancylostoma ceylanicum*

*A. ceylanicum* was maintained in Golden Syrian hamsters (Inotiv), as previously described^38,120^. Infections were established by oral gavage of 2-8 hamsters with an infective dose of ~60-100 iL3s/hamster in ~100 µL of sterile PBS. During the patency period (*i*.*e*., 14-44 days post-injection), fecal pellets were collected and mixed into coprocultures as described for the *S. stercoralis* UPD strain. Fecal-charcoal plates were kept at 23°C and iL3s were isolated from 10-11-day-old fecal-charcoal plates using a Baermann apparatus^72^.

### Maintenance of *Caenorhabditis elegans*

All *C. elegans* strains described in this paper were maintained at room temperature (~21°C) on lawns of *Escherichia coli* OP50 bacteria seeded onto 6 cm plates of 2% Nematode Growth Medium (NGM), using standard procedures^121^. Strains were passed, well-fed, for three generations prior to isolating worms for behavioral assays. Three days prior to collecting adult worms for behavioral assays, three adult worms were introduced to an OP50-seeded NGM plate. To induce the production of dauers^122^, eight adult worms were each placed onto 4-12 OP50-seeded NGM plates; dauers were collected from these plates 10-12 days later. To aid in passing social strains (which predominate in clumps around the lawn border), worm piles were initially picked to an unseeded NGM plate and allowed to spread out before individual adults were picked to a new OP50-seeded NGM plate for strain maintenance.

### Identification of nematode sGCs and phylogenetic analysis

We identified four candidate sGCs in the *S. stercoralis* genome using WormBase ParaSite v13.0 and v15.0 to perform both BlastP and TBlastN searches for homologs of proteins encoded by *Cel-gcy-31, Cel-gcy-32, Cel-gcy-33, Cel-gcy-34, Cel-gcy-35, Cel-gcy-36*, and *Cel-gcy-37*^46,62,90,123-125^. Our hits were SSTP_0000680700, SSTP_0000411800, SSTP_0000255800, and SSTP_0000482200. Phylogenetic clustering was used to classify these genes as homologs of *Cel*-sGCs. First, a MUSCLE alignment^126^ was performed on the amino acid sequences of the four candidate *Sst-*sGCs and seven *Cel*-sGCs; of note, the protein sequence for *Cel*-GCY-33 was manually annotated as described in the corresponding source data file. The alignment file was then uploaded to W-IQ-Tree^127^, using the default parameters, for maximum likelihood phylogenetic analysis. The result of this analysis (Fig. 3A) was visualized in RStudio (v2021.09.2) using the ggtree^128-130^ and treeio^131^ packages, as previously described^39^. From this analysis, we identified SSTP_0000680700 as a homolog of *Cel-gcy-35* (*Sst-gcy-35*), SSTP_0000411800 as a homolog of *Cel-gcy-36* (*Sst-gcy-36*), and both SSTP_0000255800 and SSTP_0000482200 as homologs of *Cel-gcy-37* (*Sst-gcy-37*.*1* and *Sst-gcy-37*.*2*, respectively).

To identify candidate sGCs in other nematode species, we used a WormBase ParaSite BioMart search^125^ to identify genes coding for proteins containing an H-NOX domain (InterPro ID: IPR038158)^132^. We searched for candidate sGCs in clade I species (*i*.*e*., the mammalian-infective nematodes *Ascaris lumbricoides, Trichinella spiralis*, and *Trichuris trichiura*), clade III species (*i*.*e*., *Toxocara canis* and the filarial nematodes *Loa loa, Brugia malayi*, and *Onchocerca volvulus*), clade IV species (*i*.*e*., the free-living nematode *Rhabditophanes diutinus*, the entomopathogenic nematode *Steinernema carpocapsae*, and the skin-penetrating, facultative parasitic nematodes *S. stercoralis, S. ratti, Strongyloides venezuelensis*, and *Parastrongyloides trichosuri*), and clade V species (*i*.*e*., the free-living nematodes *C. elegans, Caenorhabditis briggsae*, and *Pristionchus pacificus*; the parasitic hookworms *A. ceylanicum* and *Necator americanus*; and the passively ingested ruminant parasite *Haemonchus contortus*). Candidate sGCs were manually curated^124^; any candidate sGC hits lacking either an H-NOX domain or guanylate cyclase domain were excluded, unless the missing domain was readily apparent in the gene sequence adjacent to the hit and could be manually added. Hits with extraneous domains were truncated; where possible, relative expression levels from RNA-seq were analyzed to confirm that extraneous domains were likely transcribed independently of the sGC in question. Phylogenetic clustering analysis was performed on the predicted protein sequences, as described above. The results of this analysis (Fig. 5A) concurred with identification of SSTP_0000680700 as a homolog of *Cel-gcy-35* (*Sst-gcy-35*), SSTP_0000411800 as a homolog of *Cel-gcy-36* (*Sst-gcy-36*), and both SSTP_0000255800 and SSTP_0000482200 as homologs of *Cel-gcy-37* (*Sst-gcy-37*.*1* and *Sst-gcy-37*.*2*, respectively).

### Accession of RNA-sequencing analysis

Publicly available RNA-sequencing (RNA-seq) data for *S. stercoralis*^81,82^ was accessed using the *Strongyloides* RNA-seq Browser^80^. The expression levels (from three biological replicates) of *Sst-gcy-35, Sst-gcy-36, Sst-gcy-37*.*1*, and *Sst-gcy-37*.*2* are displayed for each life stage (Fig. 3B).

### Molecular biology and generation of transgenic *S. stercoralis* iL3s

Four candidate sGC genes (*i*.*e*., *Sst-gcy-35, Sst-gcy-36, Sst-gcy-37*.*1*, and *Sst-gcy-37*.*2*) were identified in *S. stercoralis* using protein sequence homology to *Cel*-GCY-31 through *Cel*-GCY-37, as described above. To generate a transcriptional reporter for each *Sst-*sGC gene, the promoter (*i*.*e*., a ~2000-3000 base pair (bp) region immediately upstream of transcriptional start site) was synthesized by GenScript (Piscataway, NJ, USA). Each promoter was cloned into the pSD10 vector, which expresses a *Strongyloides*-codon-optimized GFP (*strGFP*) flanked downstream by the *Sst-era-1* 3’ UTR. The 2043 bp *Sst-gcy-35* promoter was cloned into pSD10 using EcoRV-HF (New England Biolabs, R3195) and AgeI-HF (New England Biolabs, R3552) to generate pBMW14; pBMW14 was microinjected at a concentration of 50 ng/µL into *S. stercoralis* P_0_ free-living females and transgenic F_1_ iL3s were isolated from the progeny, as previously described^72^. The 2997 bp *Sst-gcy-36* promoter was cloned into pSD10 using EcoRV-HF (New England Biolabs, R3195) and AgeI-HF (New England Biolabs, R3552) to generate pBMW15; to produce transgenic F_1_iL3s, pBMW15 was microinjected at a concentration of 50 ng/µL into *S. stercoralis* P_0_ free-living females^72^. The 2999 bp *Sst-gcy-37*.*1* promoter was cloned into pSD10 using HindIII (New England Biolabs, R3104) and AgeI-HF (New England Biolabs, R3552) to generate pBMW16; to produce transgenic F_1_ iL3s, pBMW16 was microinjected at a concentration of 50 ng/µL into *S. stercoralis* P_0_ free-living females^72^. In some cases, the co-injection marker pAJ50^73^ – which drives expression of mRFPmars in the body-wall muscle – was injected at a concentration of 20 ng/µL alongside pBMW16. The 3001 bp *Sst-gcy-37*.*2* promoter was cloned into pSD10 using HindIII (New England Biolabs, R3104) and the compatible cutters XmaI (New England Biolabs, R0180) and AgeI-HF (New England Biolabs, R3552) to generate pBMW17; microinjection of pBMW17 into P_0_ free-living females^72^ at various concentrations never yielded progeny with visible expression of the transgene.

To express the light chain of tetanus toxin (TeTx) in *Sst-*URX, we generated pBMW56 by using SacI-HF (New England Biolabs, R3156) and AgeI-HF (New England Biolabs, R3552) to clone the *Sst-gcy-36* promoter from pBMW15 into pMLC212, which contains a multiple cloning site and a *strTeTx*::P2A::*strmScarlet-I*::*Sst-era-1* 3′ UTR cassette. The *strTeTx*::P2A::*strmScarlet-I* sequence was originally synthesized by GenScript (Piscataway, NJ, USA), as previously reported^42^. pBMW56 was microinjected into *S. stercoralis* P_0_free-living females at a concentration of 80 ng/µL^72^; F_1_ iL3s expressing TeTx in *Sst-*URX were selected by visible mScarlet-I expression using a Leica M165 FC fluorescence microscope. To express TeTx in *Sst-*BAG, pNB14 (*Sst-gcy-9p::strTeTx::*P2A*::strmScarlet-I::Sst-era-1* 3′ UTR) was microinjected into P_0_free-living females at a concentration of 80 ng/µL and F_1_ iL3s with visible mScarlet-I expression were selected^42,72^.

To perform ratiometric calcium imaging, we expressed *Strongyloides*-codon-optimized GCaMP7f (*strGCaMP7f*) and mScarlet-I (*strmScarlet-I*) using a polycistronic vector. The *strGCaMP7f::*P2A::*strmScarlet-I* cassette was originally synthesized by GenScript (Piscataway, NJ, USA) and used to generate pRP08. To drive expression in *Sst-*URX, we used SacI-HF (New England Biolabs, R3156) and AgeI-HF (New England Biolabs, R3552) to clone the *Sst-gcy-36* promoter from pBMW15 into pRP08, resulting in pBMW31 (*Sst-gcy-36p::strGCaMP7f::*P2A*::strmScarlet-I::Sst-era-1* 3′ UTR). To drive expression in *Sst-*BAG, we used AgeI-HF (New England Biolabs, R3552) and AvrII (New England Biolabs, R0174) to clone the *strGCaMP7f::*P2A*::strmScarlet-I* cassette from pBMW31 into pNB14, thus creating pBMW66 (*Sst-gcy-9p::strGCaMP7f::*P2A*::strmScarlet-I::Sst-era-1* 3′ UTR). Either pBMW31 or pBMW66 was microinjected into P_0_ free-living females at a concentration of 80 ng/µL and F_1_ iL3s with visible GCaMP7f and mScarlet-I expression were selected for calcium imaging^72^.

### Generation of a stable *S. stercoralis Sst-gcy-35* line

The *S. stercoralis* EAH467 *Sst-gcy-35*(*bru7*[*Sst-act-2p::strGFP::Sst-era-1* 3′ UTR]) II / *Sst-gcy-35*(*bru8*[*Sst-act-2p::strmScarlet-I::Sst-era-1* 3′ UTR]) II strain was generated using previously described methods^42,75,83^. Briefly, the CRISPR target site (5’ - GAGATTTGGGAAGCTTATGG – 3’) for *Sst-gcy-35* was identified using Geneious 9 software; this site was prioritized because of its relatively high on-target activity score calculated by the algorithm developed in Doench *et al*., 2016^133^ and its GN(17)GG motif, known to increase mutation efficiency in *C. elegans*^134^. GenScript (Piscataway, NJ, USA) synthesized the *Sr-U6p::Sst-gcy-35-sgRNA::Sr-U6* 3’ UTR cassette and cloned it into the pMLC47 backbone^83^ to generate pBMW58. To generate the homology-directed repair (HDR) plasmid (pBMW57), the 559 bp 5’ homology arm (HA) and 560 bp 3’ HA were synthesized by GenScript (Picataway, NJ, USA) and cloned into pRP12 such that the HAs flanked the *Sst-act-2p::strmScarlet-I::Sst-era-1* 3’ UTR cassette that drives body wall expression of mScarlet-I. To make an HDR plasmid with the HAs flanking an *Sst-act-2p::strGFP::Sst-era-1* 3’ UTR cassette (pBMW59) for expression of GFP in the body wall, the *strGFP* sequence from pBMW14 was cloned into pBMW57 using AgeI-HF (New England Biolabs, R3552) and AvrII (New England Biolabs, R0174). pPV540 was used to drive expression of a *Strongyloides*-codon-optimized *Cas9* gene, as previously described^83,135^.

To create the *S. stercoralis Sst-gcy-35* stable line, 107 P_0_ free-living females were microinjected with a mixture of pBMW59 (80 ng/µL), pBMW58 (80 ng/µL), and pPV540 (50 ng/µL) and 128 P_0_ free-living females were microinjected with a mixture of pBMW57 (80 ng/µL), pBMW58 (80 ng/µL), and pPV540 (50 ng/µL) over the course of two days. The injected free-living females were added to fecal-charcoal plates, along with non-injected free-living males; 5-6 days post-injection, iL3s were isolated using a Baermann apparatus and progeny expressing either GFP (156 iL3s) or mScarlet-I (131 iL3s) throughout their body wall were selected for infections.

The 287 iL3s were activated *in vitro* for ~40 hours, using established methods^42,75,119^. All 287 worms were pooled, washed 3X in sterile PBS, and introduced into a Mongolian gerbil in 200 µL of sterile PBS by oral gavage.

During the patency period (days 14-44 post-infection), feces were collected from the inoculated gerbil and F_2_/F_3_ iL3s contained therein were screened for dual expression of GFP and mScarlet-I. We considered iL3s expressing both GFP and mScarlet to have a presumed homozygous knockout of *Sst-gcy-35* (*i*.*e*., the *Sst-act-2p::strGFP::Sst-era-1* 3′ UTR cassette integrated into one chromosome and the *Sst-act-2p::strmScarlet-I::Sst-era-1* 3′ UTR cassette integrated into the other), and confirmed that dual-color iL3s were *Sst-gcy-35* homozygous knockouts by single-worm genotyping (described below). To propagate the line, dual-color iL3s were picked, activated *in vitro*, and introduced to a second round of gerbils by oral gavage. Continued maintenance of this line was performed by serial passage through Mongolian gerbils using oral gavage, as described above.

### Single-worm genotyping

Single-worm genotyping was performed as previously described^38,39,41,42,83^, using primer sets (Table S1) for the wild-type and mutated *Sst-gcy-35* loci. Individual iL3s were transferred to a PCR tube in 6 µL of nematode lysis buffer (50 mM KCl, 10 mM Tris pH 8, 2.5 mM MgCl_2_, 0.45% Nonidet-P40, 0.45% Tween-20, 0.01% gelatin in ddH_2_O) supplemented with ~0.12 μg/μL Proteinase-K and ~1.7% 2-mercaptoethanol) and transferred to -80°C for at least one day. Digestion of the worm was performed using a thermocycler set to 65°C (2 h), 95°C (15 min), 10°C (hold). To perform genotyping, 1.8 µL of each digest was dispensed into one of three 50 µL PCR reactions (*i*.*e*., wild-type, 3’ integration, and control). PCR reactions were performed using Platinum™ *Taq* DNA Polymerase, High Fidelity (ThermoFisher, 11304011) and the following thermocycler conditions: 2 min denaturation at 94°C; 35 cycles of 94°C (15 s), 53°C (30 s), and 68°C (1 min); 5 min final extension at 68°C; 10°C hold.

### Gene map of *Sst-gcy-35*

The gene map for *Sst-gcy-35* (Fig. 3C) was created using Exon-Intron Graphic Maker (Version 4, www.wormweb.org). The intron and exon boundaries for SSTP_0000680700 (*Sst-gcy-35*) were accessed using WormBase ParaSite^123,125^. The CRISPR target site was identified using Geneious 9 software, as described above.

### Preparation of worms for acute O_2_ shift assays

Wild-type *S. stercoralis* iL3s and wild-type *S. ratti* iL3s were isolated from 7-10-day-old fecal-charcoal plates using a Baermann apparatus for ~90 min. Wild-type *A. ceylanicum* iL3s were isolated from 10-11-day-old fecal-charcoal plates using a Baermann apparatus for ~90 min. iL3s from all three species were washed 2-3X in BU saline^136^ and transferred to a shallow dish of BU saline until used for assays^136^.

Using a ~90 min Baermann apparatus, transgenic F_1_ *S. stercoralis* iL3s were isolated from fecal-charcoal plates 5-8 days post-microinjection of P_0_ FLFs. *S. stercoralis Sst-gcy-35* iL3s were isolated from 7-10-day-old fecal-charcoal plates using a Baermann apparatus for ~90 min. Worms were deposited onto 6 cm 2% NGM plates^121^ seeded with *E. coli* OP50 and screened for transgene expression using a Leica M165 FC fluorescence microscope. Transgene-positive iL3s were picked into a shallow dish of BU saline using a paintbrush, until the start of assays.

*S. stercoralis* FLFs and FLMs were isolated by a ~90 min Baermann apparatus from fecal-charcoal plates that were incubated at 25°C for 1 day. Free-living adults were washed 2-3X in BU saline and then deposited dropwise onto the surface of a 10 cm 2% NGM. Worms were kept on the 2% NGM plate – without food – for a minimum of 3 h prior to the assay start. Worms were picked directly from the 2% NGM plate via paintbrush for assays. Assayed *S. stercoralis Sst-gcy-35* FLFs were verified *post hoc* to be dual-color (red/green) homozygous knockouts using a Leica M165 FC fluorescence microscope.

*C. elegans* adults were prepared using standard methods^121^, with modifications. Three days prior to assays, three *C. elegans* adults were transferred to each of 8-15 6 cm 2% NGM plates seeded with *E. coli* OP50. On the assay day, worms were washed from the plate surfaces in M9 saline, collected in a 15 mL conical tube, and washed 2-3X in M9 saline. *C. elegans* adults were then deposited dropwise onto the surface of a 10 cm 2% NGM. Worms were kept on the 2% NGM plate – without food – for a minimum of 3 h prior to the assay start. For assays, adult worms with fewer than 10 eggs were picked via paintbrush from the 2% NGM plate.

*C. elegans* dauers were prepared using standard methods^122^. *C. elegans* dauers were isolated from 6 cm 2% NGM plates that had each been inoculated with 8 *C. elegans* adults 10-12 days prior to assays. These plates were seeded with *E. coli* OP50 when adult worms were added; 10-12 days later, all OP50 had been consumed. On the assay day, worms were washed from the plate surfaces in dH_2_O and collected in a 15 mL conical tube. Worms were pelleted and then treated with 5 mL of 1% SDS for 15 min (with gentle agitation) to kill non-dauer stages. Worms were then washed 3X in dH_2_O to remove residual SDS and then transferred to a shallow dish of dH_2_O until the start of assays. Within 1 h of an assay, worms were dispensed dropwise onto the surface of a 10 cm 2% NGM plate, such that live dauers could be easily isolated from worm carcasses. Worms were picked directly from the 2% NGM plate via paintbrush for assays.

### Acute O_2_ shift assays and behavioral tracking

Acute O_2_ shift assays were performed by exposing worms to different concentrations of O_2_ while crawling freely on a 14 cm 2% NGM plate. Residual moisture was removed from the agar surface by placing the uncovered plates in a fume hood for 30-120 min, depending on the relative humidity. Dried plates were then left on the bench top to acclimate to room temperature (~21°C). For assays with *S. stercoralis* iL3s, *A. ceylanicum* iL3s, and *S. ratti* iL3s, worms were first briefly rinsed in sterile dH_2_O and then transferred to the plate surface in 2 µL droplets. For assays with *S. stercoralis* FLFs, *S. stercoralis* FLMs, *C. elegans* dauers, and *C. elegans* adults, 2 µL droplets of sterile dH_2_O were pre-loaded onto the agar surface and worms were transferred into each droplet via a paintbrush (Symphony nail liner brush, UPC #053742756612, Amazon.com).

Once the worm-containing droplets dried, a chamber lid (gift from Manuel Zimmer) with a 6 cm viewing window was placed on the plate surface and secured with a weight around the perimeter, as previously described^42,49,50^. Flexible PVC tubing connected to the gas outflow port of the chamber lid was fed into a small water reservoir, where the presence of bubbles indicated an adequate seal throughout the assay. On the opposite side of the chamber lid, flexible PVC tubing supplied gas mixtures (21% O_2_/balance N_2_, 7% O_2_/balance N_2_) from certified pre-mixed tanks (AirGas) via the gas inflow port. Gas inflow from two tanks was regulated by valves controlled by a ValveBank TTL pulse generator (AutoMate Scientific) and fed into the single gas inflow port of the chamber lid using a Y-shaped connector. Gases were introduced at a flow rate of 600 mL/min (controlled by flow meters; VWR, FR2A138VVT-VW). To ensure rapid switching of the O_2_ level within the chamber, we placed a Self-Adhesive Oxygen Sensor Spot (PreSens, SP-PSt3-SA23-D5-YOP-US) on the inner side of the chamber lid’s viewing window and measured the O_2_ level within the chamber using a Polymer Optical Fiber (PreSens, POF-L2.5-1SMA) connected to a Single Channel Fiber Optic Oxygen Transmitter (PreSens, OXY-SMA-1). O_2_ levels during acute shifts were captured using PreSens Measurement Studio 2 software (Fig. S1B).

For air controls (Fig. S1C-H), worms were allowed to acclimate to the plate surface while being exposed to 21% O_2_ for 1 min. We then video-recorded worms for 2 min, while exposing them to a 40 s pulse of 21% O_2_ (from tank 1), followed by a 40 s pulse of 21% O_2_ (from tank 2), and then a 40 s pulse of 21% O_2_ (from tank 1); switching tanks allowed us to control for mechanosensory responses to pressure changes at the time of valve switching. To first expose worms to an upshift in O_2_ concentration followed by a downshift (Fig. S2), worms were allowed to acclimate to the plate surface while being exposed to 7% O_2_ for 1 min. We then video-recorded worms for 2 min, while exposing them to a 40 s pulse of 7% O2, followed by a 40 s pulse of 21% O2, and then a 40 s pulse of 7% O2. To first expose worms to a downshift in O2 level followed by an upshift (Fig. 1-4, 6, S1-S2, S4-S5, and S8), worms were allowed to acclimate to the plate surface while being exposed to 21% O_2_ for 1 min. We then video-recorded worms for 2 min, while exposing them to a 40 s pulse of 21% O_2_, followed by a 40 s pulse of 7% O_2_, and then a 40 s pulse of 21% O_2_.

Worms were video-recorded using a Leica S9D microscope (equipped with a 0.5x supplementary lens, a 300 mm M-series Focus drive, and a TL3000 ergo transmitted light base) with an attached Basler Ace 20-megapixel acA5472-17μm camera (mounted on a 0.63x Leica adapter lens). Images were captured at 10 frames/s using pylon Viewer software (Basler, Exton, PA, USA). Captured images for each assay were converted to a .AVI file using Fiji^137^ and movement parameters were quantified using WormLab (MBF Bioscience, Williston, VT, USA) automated tracking software. For each worm, the smoothed speed was measured throughout the 2 min assay; the moving average speed over 3 s was smoothed using locally weighted polynomial regression. We then determined the mean smoothed speed, forward time, reverse time, and/or idle time of each worm during the first and last 10 s of each 40 s gas pulse. A worm was considered idle if traveling at an absolute speed of <30 µm/s for at least 1 s; a worm was called as reversing if traveling tail-first at a minimum absolute speed of at least 30 µm/s for at least 1 s. To quantify rapid, transient pauses provoked by a change in the ambient O_2_ concentration (Fig. 2E-F, 4H-I, and S8B), we compared the minimum speed of worms in the 5 s preceding an O_2_ level shift with the minimum speed of worms in the 5 s following the O_2_ level shift. Worms were excluded from analysis if tracking could not be performed for at least 95% of the recording period; worms were also excluded if the track was incomplete (missing frames) in a binning region used for statistical analysis. Worms often left the field of view mid-assay and were subsequently excluded. Other exclusion criteria include: 1) collisions with other worms resulting in track breaks of >5 s and/or an inability to distinguish the touching worms; 2) inaccurate calls by the automated tracking software following manual curation of the track; and 3) poor track quality and/or an inability to maintain a track due to debris on the plate, optic disturbances, and/or worm movement into condensed/coiled body shapes.

### Thermal stimulation assays and speed tracking

Wild-type and *Sst-gcy-35 S. stercoralis* iL3s were isolated from fecal-charcoal plates as described above. iL3s were stored in a thin layer of BU^136^ in a watch glass until the assay start. Assays were performed on 10 cm 2% NGM plates. Residual moisture was removed from the agar surface by placing the uncovered plates in a fume hood for 90 min. Dried plates were then either left on the bench top to acclimate to room temperature (~21°C) or transferred to a 37°C incubator to pre-heat. To perform assays at room temperature, iL3s were briefly rinsed in sterile dH_2_O and then deposited on the plate surface in 2 µL droplets. Once the droplets dried, worms were video-recorded for 45 s using a Leica S9D microscope (equipped with a 0.5x supplementary lens, a 300 mm M-series Focus drive, and a TL3000 ergo transmitted light base) with an attached Basler Ace 20-megapixel acA5472-17μm camera (mounted on a 0.63x Leica adapter lens). Images were captured at 10 frames/s using pylon Viewer software (Basler, Exton, PA, USA). Assays performed at host body temperature (34-37°C)^38^ were performed identically, with two modifications: 1) pre-heated plates were placed on the Lecia S9D microscope stage equipped with a Leica MATS heating system set to 57.2°C and 2) the plate surface temperature was measured pre- and post-assay using an infrared laser thermometer (TackLife, IT-T02) to ensure that the plate surface temperature did not fall outside of the 34-37°C range.

Captured images for each assay were converted to a .AVI file using Fiji^137^. The mean smoothed speed of each iL3 during the initial 30 s of the recording was quantified using WormLab (MBF Bioscience, Williston, VT, USA) automated tracking software. The moving average speed over 3 s was smoothed using locally weighted polynomial regression.

### Fluorescence microscopy

Epifluorescence and differential interference contrast (DIC) imaging were performed as previously described^42,75^, using an inverted Zeiss AxioObserver A2 microscope equipped with a Plan-APOCHROMAT 20X objective lens, a Colibri 7 (Zeiss) for LED fluorescence illumination, a 38 HE filter set for GFP (BP470/40, FT495, BP 525/50), a 63 HE filter set for mScarlet-I (BP572/25, FT590, BP629/62), a Hamamatsu ORCA-Flash4.0 camera, and Zen 3.3 (blue edition; Zeiss) software. Heated Noble agar (5% in BU saline) was deposited dropwise onto a microscope slide and flattened to form a pad. iL3s were paralyzed using 50-100 μM levamisole (dissolved in BU saline), placed onto the agar pad, and secured with a coverslip. Images displayed in Fig. S6E were captured as a snap of a single plane; all other images were captured as z-stacks. Fiji^137^ was used to pseudo-color all images and to convert z-stacks to maximum intensity projection images.

### Calcium imaging

Ratiometric calcium imaging was performed using transgenic *S. stercoralis* iL3s expressing both GCaMP7f^95^ and mScarlet-I in neurons of interest. One day prior to imaging, iL3s were isolated from fecal-charcoal plates using a Baermann apparatus and paralyzed on the surface of a 10 cm 2% NGM plate using 2% nicotine (dissolved in BU saline). iL3s with neuronal expression of GCaMP7f and mScarlet-I were transferred to a shallow dish of BU saline and kept overnight^136^.

Coverslips (48 × 60 mm, Brain Research Laboratories, 4860-1D) seeded with a small droplet of 2% Noble agar (dissolved in BU saline) were also prepared one day prior to imaging and left to dry in a covered tray overnight. Immediately before the start of imaging, a single iL3 was transferred in a 2 µL drop of BU saline into the center of the dried agar. A thin strip of Whatman paper was placed near the edge of the BU saline droplet and used to wick away excess liquid. Once the BU saline droplet fully dissipated, the iL3 adhered to the dried agar, thus immobilizing the animal for imaging. A self-adhesive perfusion chamber (20 mm diam. x 0.9 mm depth, Grace Bio-Labs CoverWell™, Millipore Sigma, GBL622101) was then placed over each worm and pressed securely to the coverslip surface to create an airtight seal. Gas flowed into the perfusion chamber via a 1.5 mm port, where flexible PVC tubing was connected using a self-adhesive press fit tubing connector (Grace Bio-Labs, Millipore Sigma, GBL460003). A self-adhesive press fit tubing connector was also used to connect the 1.5 mm gas outflow port (located 180° opposite the inflow port) to flexible PVC tubing, which was fed into small water reservoir; the presence of bubbles in the outflow tract water reservoir confirmed the perfusion chamber’s seal to the coverslip throughout the duration of imaging.

Gas mixtures (21% O_2_/balance N_2_, 7% O_2_/balance N_2_, or 15% CO_2_/21% O_2_/balance N_2_) were delivered from certified pre-mixed tanks (AirGas) using flexible PVC tubing at a flow rate of 28.2 mL/min (using a Supelco Rotameter with a Carboloy™ float, Millipore Sigma, 23324). To reduce the risk of worm desiccation during imaging, gases were humidified using 250 mL borosilicate Dreschel gas washing bottles (Eisco, VWR, 470344-412) containing dH_2_O. Gas inflow from each tank was modulated by valves controlled by a ValveBank TTL pulse generator (AutoMate Scientific) and fed into the single gas inflow port of the perfusion chamber using a Y-shaped connector; backflow around the Y-shaped connector was blocked using one-way check valves (ANPTGHT, Amazon, B091T7MQXY). For air controls (Fig. 7, Fig. S10), iL3s were exposed to a 40 s pulse of 21% O_2_ (from tank 1), followed by a 40 s pulse of 21% O_2_(from tank 2), and then an 80 s pulse of 21% O_2_ (from tank 1); switching tanks allowed us to control for mechanosensory responses to pressure changes at the time of valve switching. For low O_2_ exposure (Fig. 7), iL3s were exposed to a 40 s pulse of 21% O_2_, followed by a 40 s pulse of 7% O_2_, and then an 80 s pulse of 21% O_2_. For high CO_2_ exposure (Fig. S10), iL3s were exposed to a 40 s pulse of 21% O_2_, followed by a 40 s pulse of 15% CO_2_/21% O_2_, and then an 80 s pulse of 21% O_2_.

Imaging was performed with an upright Zeiss AxioImager A2 microscope equipped with a 40x objective (EC Plan-Neofluar 40x/0.75 ∞/0.17; Zeiss), a Colibri 7 (Zeiss) for LED fluorescence illumination, a 79 HE ms filter set (BP 470/28 + BP 556/25, DBS 490+575 + FT 565, BP 512/30 + BP 630/98; Zeiss), a Hamamatsu W-View Gemini beam splitter with a GFP/mRFP dual camera filter set, and a Hamamatsu ORCA-Flash4.0 camera for simultaneous acquisition of GCaMP7f and mScarlet-I images. Images were captured at 2 frames/s using Zen 3.9 (blue edition; Zeiss) software.

Images were processed using Zen 3.9 (blue edition; Zeiss) and Microsoft Excel. First, simultaneously acquired images of GCaMP7f and mScarlet-I signal were overlayed, and a region of interest (ROI) was drawn around the cell body of the imaged neuron. This ROI was confirmed to contain the neuron cell body in every frame of the captured images, and the average intensities of the GCaMP7f and mScarlet signals were separately measured. A second ROI of equal area was drawn in the background region. For GCaMP7f, the average signal intensity was measured in the background ROI and subtracted from the average signal intensity in the neuron cell body ROI; the same background subtraction was performed for mScarlet-I. For each frame, the ratio of GCaMP7f to mScarlet-I signal was calculated. To calculate the change in fluorescence intensity from baseline (%ΔR/R_0_), the ratio of GCaMP7f to mScarlet-I signal in each frame was compared to the average GCaMP7f to mScarlet-I signal in the final 10 s of the first 40 s gas pulse; this 10 s baseline period is displayed at t = 0-10 s in all graphs. All GCaMP7f to mScarlet-I signal ratios were subject to baseline correction. To generate heatmaps displaying neuronal activity for individual worms, responses were ordered by using Heatmapper^138^ to perform hierarchical clustering, using the average linkage clustering method and Euclidean distance measurement method.

### *In vitro* activation assays

*In vitro* activation assays were performed essentially as previously described^41,42,105,106^, but with the ambient O_2_ concentration fixed at either 21% or 7% O_2_. First, iL3s were isolated from fecal-charcoal plates using a Baermann apparatus. They were then washed 2-3x in BU saline^136^, and resuspended in 10 mL BU saline containing 100 µL 100x penicillin-streptomycin (10,000 u/mL, Gibco 15140-122), 100 µL 100x amphotericin B (250 µg/mL, Gibco 15290-018), and 5 µL 2000x tetracycline hydrochloride (10 mg/mL, Sigma-Aldrich T7660-5G). Depending on the total number of worms collected from each Baermann, worms were sometimes axenized in 5 mL of BU saline containing 50 µL 100x penicillin-streptomycin, 50 µL 100x amphotericin B, and 2.5 µL 2000x tetracycline hydrochloride. iL3s were axenized for 3 h at room temperature in the dark and then centrifuged at 4400 rpm for 1-2 min. The supernatant was removed and the iL3s were diluted to a density of ~40-50 iL3s/µL.

A 96-well plate was prepared that had 110 µL DMEM (Gibco 11995-065), along with penicillin-streptomycin, amphotericin, and tetracycline hydrochloride at the concentrations described above, in each of 12 experimental wells (Fig. 8A). The outer wells were filled with ddH_2_O to retain moisture. The plate was preincubated at 37°C with 5% CO_2_ at either 7% O_2_ or 21% O_2_ for 1-3 h. iL3s were then added to the DMEM-filled wells by pipetting 2-5 µL of the iL3 solution into each well such that each well contained ~100-200 iL3s. For each O_2_ concentration, 6 out of the 12 wells of the plate contained wild-type iL3s, whereas the remaining 6 wells contained *Sst-gcy-35* iL3s. The plate was then incubated at 37°C with 5% CO_2_ and either 7% O_2_ or 21% O_2_ for 21 h to allow the worms to activate. To quantify activation rates, 2.5 µL of 1 mg/mL Alexa Fluor− 594 NHS Ester (ThermoFisher A20004, 1 mg/mL in N,N-dimethylformamide) was added to each well and the plate was incubated under the same conditions for an additional 3 h. For each genotype and O_2_ concentration, iL3s were then pipetted out of the wells into 1.5 mL microcentrifuge tubes and then centrifuged at 13,000 rpm for 1-2 min; the supernatant was then removed. Worms were then washed 4 times in BU saline. Worms were resuspended in ~100-200 µL BU saline and transferred in 5-10 µL droplets to a 2% NGM agar plate. To paralyze worms, 2% nicotine was added to each droplet. After the droplets dried out, the plate with the worms was observed under a Leica M165 FC fluorescence dissecting microscope. iL3s were categorized as activated if they contained red fluorescence in their pharynx. For each trial, at least 200 iL3s were scored per condition.

### Quantification and statistical analysis

Statistical testing was performed using GraphPad Prism v10.6.0 and v10.6.1. Descriptions of specific statistical analyses are included in the associated figure caption. Data were checked for normality in Prism and non-parametric tests were used for non-parametrically distributed data. Power analysis was performed using G*Power 3.1.9.6.

## Supporting information

Walsh et al Supplemental Data

## DATA AVAILABILITY

All data necessary for the conclusions described in this study are included with this article. Source data are provided with this paper. All raw data for this study are available from GitHub (https://github.com/HallemLab/Walsh_et_al_2026). Genetically modified nematodes used in this study are available upon request.

## CODE AVAILABILITY

All code used in this study is available from GitHub (https://github.com/HallemLab/Walsh_et_al_2026).

## ACKNOWLEDGMENTS

We thank Michelle Castelletto, Ruhi Patel, and Julian Wagner for thoughtful comments on the manuscript. The illustrations shown in Fig. 1A, Fig. S3A-B, Fig. 2A, Fig. 2C, Fig. 3D, Fig. 3F, Fig. S6A, and Fig. 8A were created with BioRender.com. This work was supported by NIH F30AI179222, UCLA-Caltech MSTP Training Grant NIH T32GM152342, and UCLA Molecular Biology Institute Whitcome Pre-Doctoral Fellowship in Molecular Biology to B.W.; NIH F32AI147617 and NIH T32AI007323 (PI: P. Johnson) to N.B.; NIH T32GM145388 and NIH T32AI007323 to G.B.; and NIH R01DC017959 to E.A.H.

## AUTHOR CONTRIBUTIONS

B.W. and E.A.H. conceived the study. B.W., N.B., G.B., and E.A.H. designed experiments. B.W., N.B., and G.B. performed experiments. B.W. and E.A.H. wrote the manuscript and prepared figures. All authors reviewed and provided feedback on the manuscript.

