## Supplementary material for "Neuronally sensed oxygen drives behavior and development in human-infective, skin-penetrating nematodes": Walsh et al Supplemental Data

### SUPPLEMENTAL FIGURES

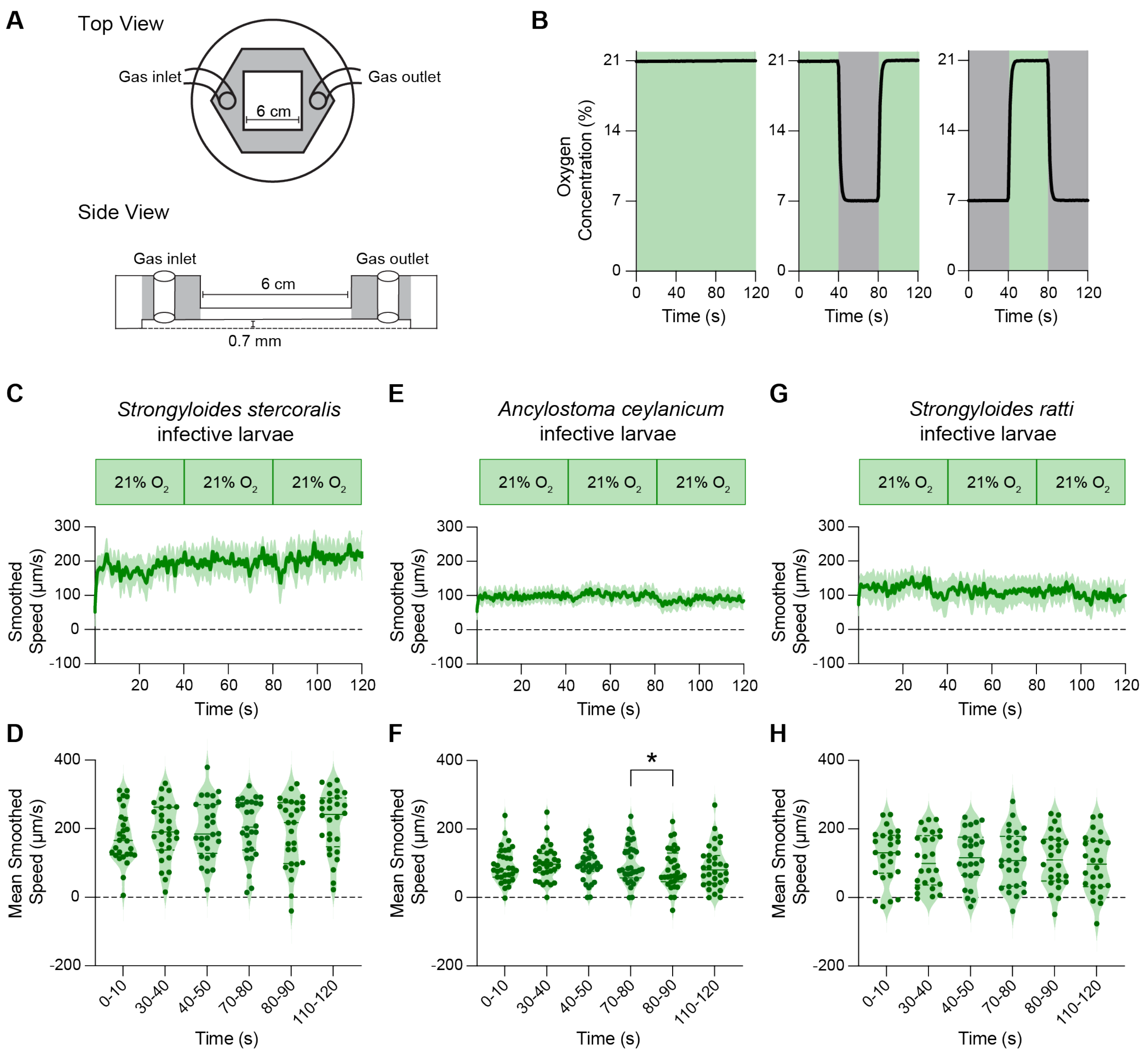


**Fig. S1. Worms are exposed to rapid shifts in $O_2$ concentration using a plate-based behavioral arena. A.** Schematic of the chamber lid[1-3] used to deliver rapid shifts in $O_2$ concentration to worms crawling on an agar plate. The plate lid, with a 6 x 6 cm optically clear viewing window (top view) seals around its outer perimeter to an agar plate. The distance between the agar plate surface and the bottom of the viewing window is 0.7 mm (side view), allowing enough space for worms to crawl freely while minimizing turbulence. **B.** The small volume of the chamber lid allows for rapid shifts in $O_2$ levels, as measured by a Single Channel Fiber Optic Oxygen Transmitter (PreSens, OXY-SMA-1). When three successive 40 s pulses of 21% $O_2$ enter the chamber lid, the $O_2$ concentration is maintained at a steady level of 21%. When the 40 s pulses shift between 21% (green) and 7% (gray), the chamber quickly floods with the new gas mixture, resulting in acute shifts in the $O_2$ level. **C-H.** In an air control, iL3s are exposed to three successive 40 s pulses of 21% $O_2$ to account for any effects of mechanical stimulation during shifts. *S. stercoralis* iL3s (**C-D**) and *S. ratti* iL3s (**G-H**) move at consistent speeds for the entire assay duration. *A. ceylanicum* iL3s also tend to move at a consistent speed throughout the assay but appear slightly more sensitive to the mechanical stimulus between successive pulses (**E-F**). In C, E, and G, bold lines show mean smoothed speed, and shading represents the 95% confidence interval. In D, F, and H, dots represent individual worms, solid lines show medians, and dotted lines show quartiles. Negative values indicate reverse movement. **p*<0.05, Friedman test with Dunn's multiple comparisons post-test; only significant, adjacent comparisons are shown. n = 26-32 iL3s per species.

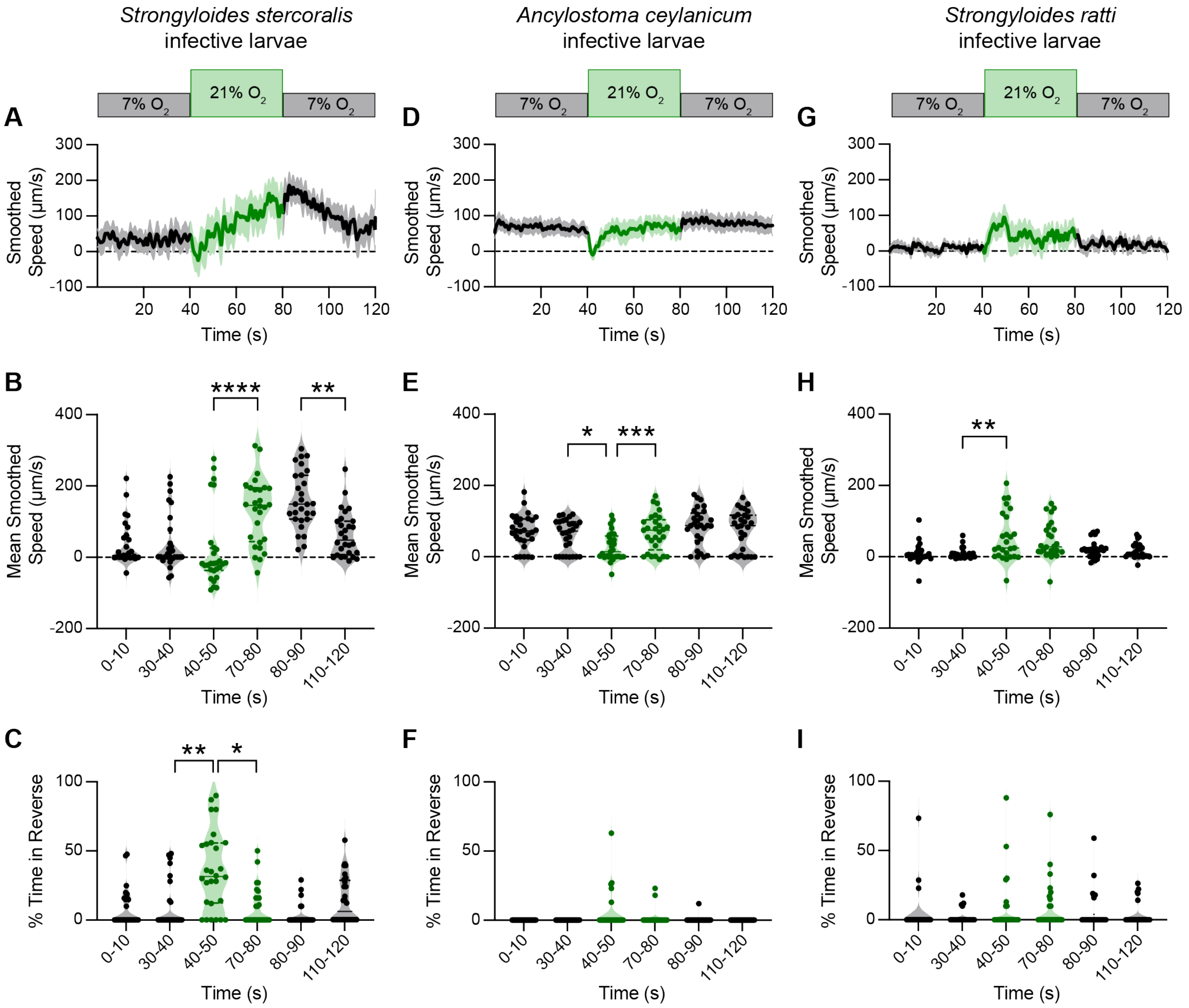


**Fig. S2. Parasitic iL3s exhibit $O_2$-evoked behaviors when first exposed to an $O_2$ downshift. A-C.** Response of *S. stercoralis* iL3s to acute $O_2$ shifts. **A.** Graph of smoothed speed. Bold line shows mean smoothed speed; shading shows 95% confidence interval. Negative values indicate reverse movement. **B.** Violin plot of mean smoothed speed during the first and last 10 s of each gas pulse. Negative values indicate reverse movement. **C.** Violin plot showing the percentage of time spent in reverse during the first and last 10 s of each gas pulse. **D-F.** Response of *A. ceylanicum* iL3s to acute $O_2$ shifts. Graphs are as described in A-C. **G-I.** Response of *S. ratti* iL3s to acute $O_2$ shifts. Graphs are as described in A-C. For A-I, iL3s were exposed to a 40 s pulse of 7% $O_2$ (black), followed by a 40 s pulse of 21% $O_2$ (green), followed by a 40 s pulse of 7% $O_2$ (black). For violin plots, dots represent individual worms, solid lines show medians, and dotted lines show quartiles. **p*<0.05, ***p*<0.01, ****p*<0.001, *****p*<0.0001, Friedman test with Dunn's post-test; only significant, adjacent comparisons are displayed. n = 26-29 iL3s per species.

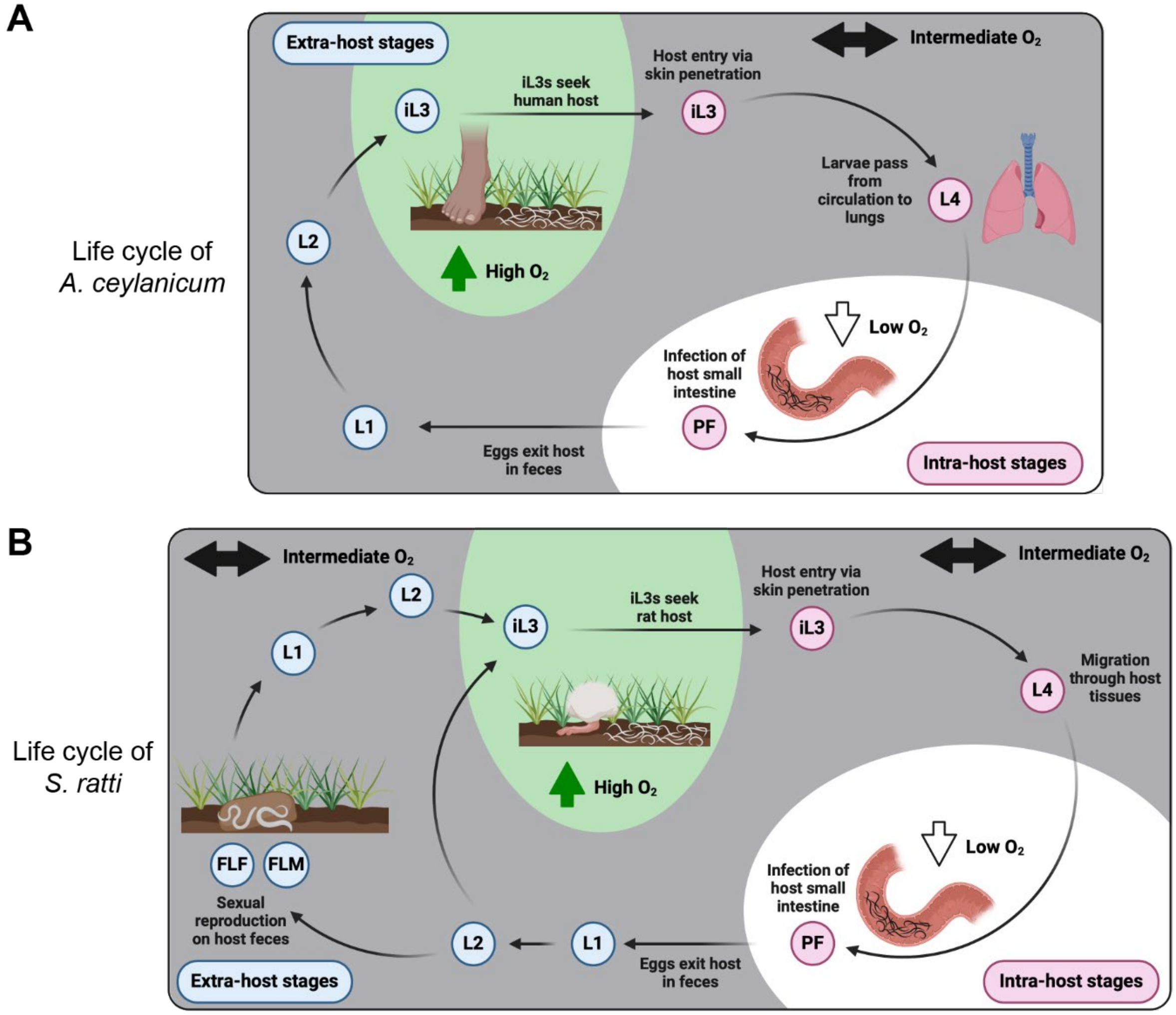


**Fig. S3. Parasitic nematodes encounter a range of $O_2$ levels throughout their life cycles. A.** The life cycle of *A. ceylanicum*[4]. iL3s reside at the soil surface (~21% $O_2$[5]), until they locate and enter a human host via skin penetration. iL3s experience an immediate drop in $O_2$ level following skin-penetration[6]. iL3s develop into infective fourth-stage larvae (L4s) that pass through the lungs (~13.5% $O_2$[6]), where they are coughed up and swallowed into the digestive tract. Parasitic females (PFs) lay eggs in the small intestine (near-anaerobic environment in lumen[6,7]), which pass from the host in feces. Eggs develop into first-stage larvae (L1s), then second-stage larvae (L2s), and finally into iL3s, completing the cycle. **B.** The life cycle of *S. ratti*[8]. iL3s seek a rat host at the soil surface (21% O2[5]). iL3s encounter intermediate, subatmospheric levels of $O_2$ upon skin penetration and as they develop into L4s while migrating through host tissues. Ultimately, PFs take up residence in the small intestine, where the $O_2$ levels are nearly anaerobic[9,10]. Eggs laid by the PFs exit the host in feces and develop into post-parasitic L1s and subsequently into post-parasitic L2s. Upon host exit, L1s and L2s (as well as other extra-host stages) experience intermediate $O_2$ levels while developing on host feces[11]. L2s can either develop directly into iL3s or develop into a single generation of free-living females (FLFs) and free-living males (FLMs) that reproduce sexually to produce post-free-living L1s. All the progeny of FLFs and FLMs eventually develop from post-free-living L2s into iL3s. For A-B, pink circles depict intra-host life stages; blue circles depict extra-host life stages. $O_2$ levels are noted by background color (high $O_2$ in green, intermediate $O_2$ in gray, and low $O_2$ in white).

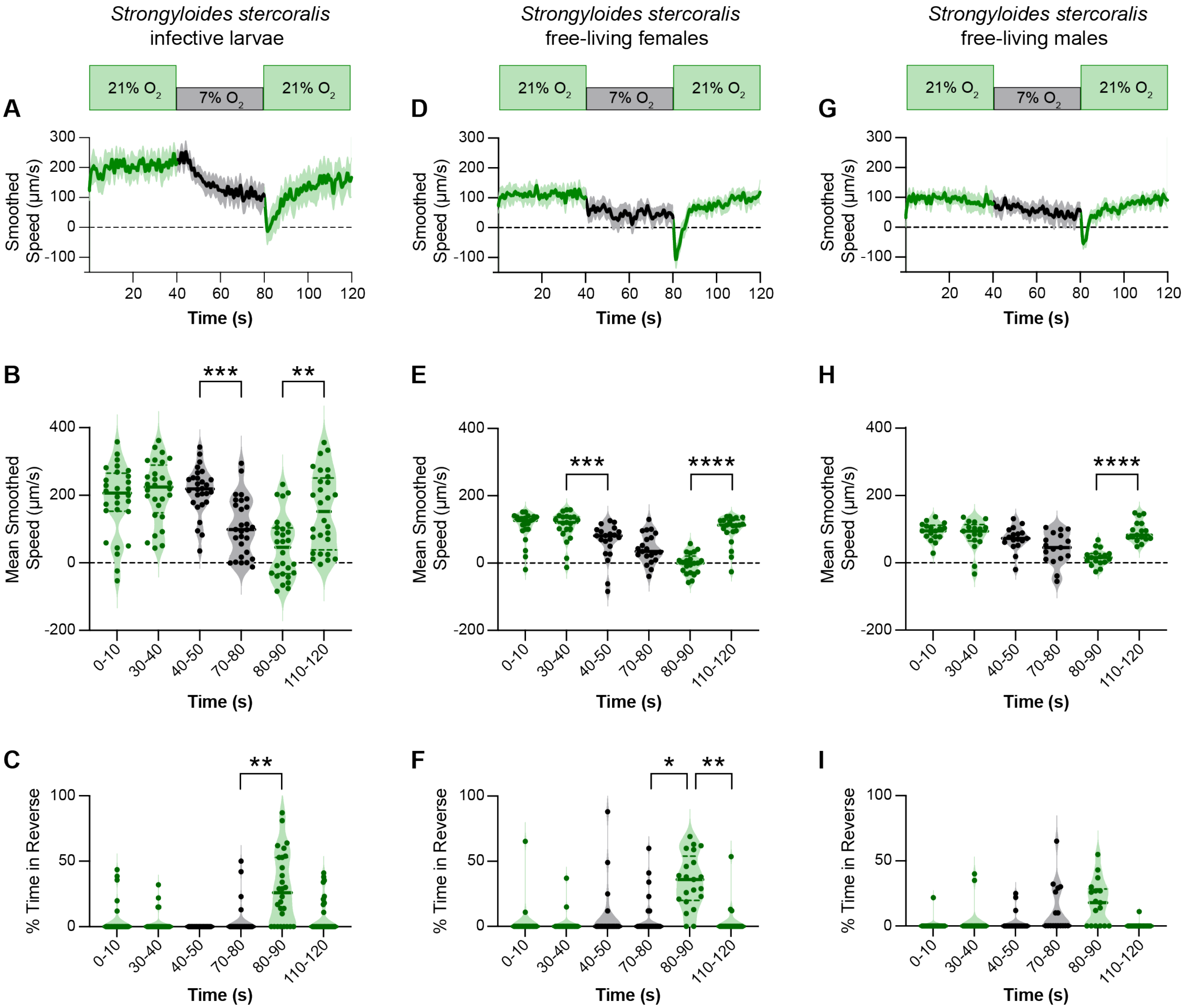


**Fig. S4.** ***S. stercoralis*** **free-living adults display $O_2$-evoked behavior. A-C.** Response of *S. stercoralis* iL3s to acute $O_2$ shifts. **A.** Graph of smoothed speed. Bold line shows mean smoothed speed; shading shows 95% confidence interval. Negative values indicate reverse movement. **B.** Violin plot of mean smoothed speed during the first and last 10 s of each gas pulse. Negative values indicate reverse movement. **C.** Violin plot showing the percentage of time spent in reverse during the first and last 10 s of each gas pulse. **D-F.** Response of *S. stercoralis* FLFs to acute $O_2$ shifts. Graphs are as described in A-C. **G-I.** Response of *S. stercoralis* FLMs to acute $O_2$ shifts. Graphs are as described in A-C. For A-I, worms were exposed to a 40 s pulse of 21% $O_2$ (green), followed by a 40 s pulse of 7% $O_2$ (black), followed by a 40 s pulse of 21% $O_2$ (green). For violin plots, dots represent individual worms, solid lines show medians, and dotted lines show quartiles. **p*<0.05, ***p*<0.01, ****p*<0.001, *****p*<0.0001, Friedman test with Dunn's post-test; only significant, adjacent comparisons are displayed. n = 17-28 worms per life stage. Data in A-C are from Fig. 1.

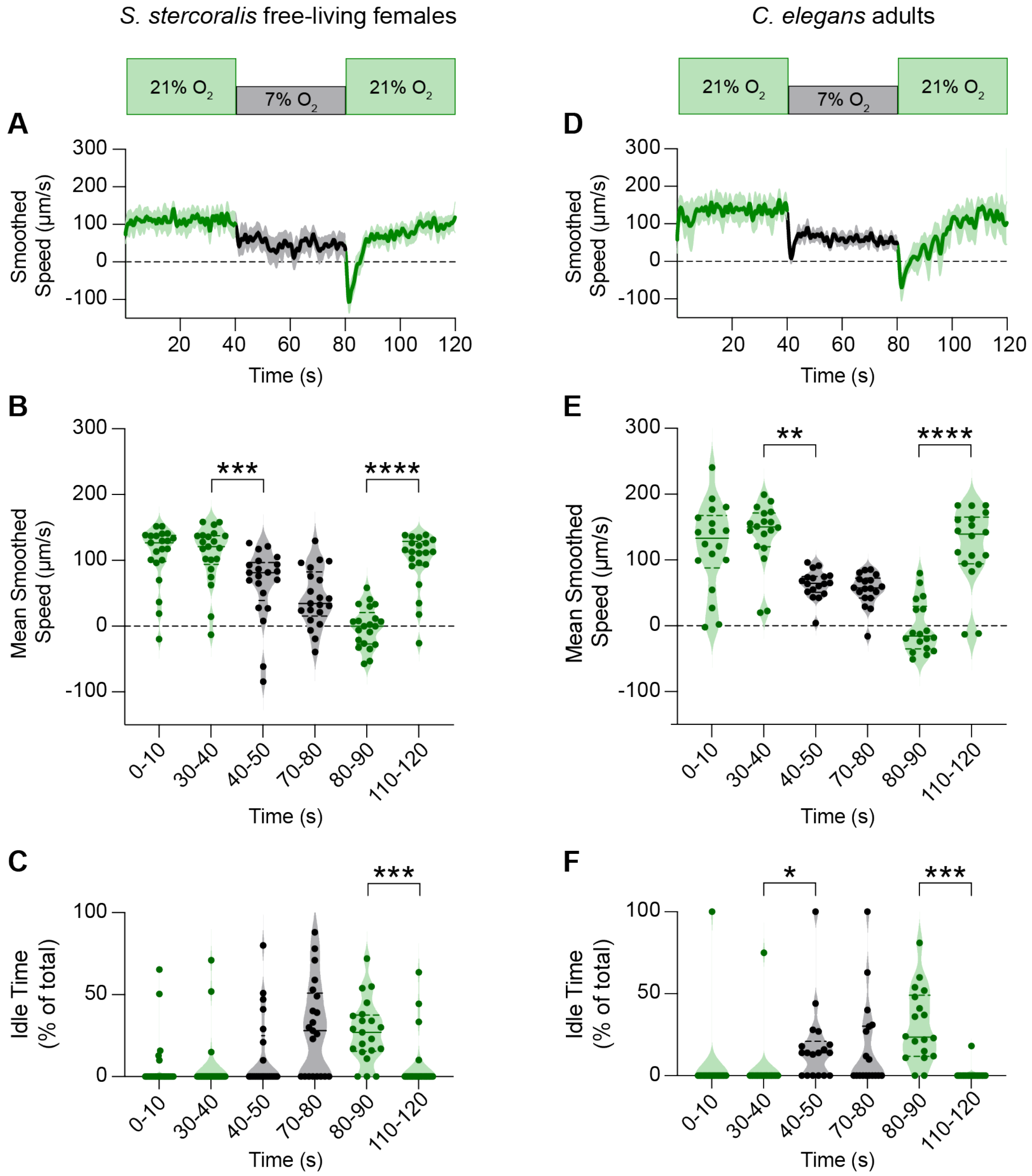


**Fig. S5.** ***C. elegans*** **adults exhibit a pause response not seen in** ***S. stercoralis*** **FLFs upon encountering an $O_2$ downshift. A-C.** Response of *S. stercoralis* FLFs to acute $O_2$ shifts. **A.** Graph of smoothed speed. Bold line shows mean smoothed speed; shading shows 95% confidence interval. Negative values indicate reverse movement. **B.** Violin plot of mean smoothed speed during the first and last 10 s of each gas pulse. Negative values indicate reverse movement. **C.** Violin plot showing the percentage of time spent idle during the first and last 10 s of each gas pulse. **D-F.** Response of *C. elegans* HW adults to acute $O_2$ shifts. Graphs are as described in A-C. For A-F, worms were exposed to a 40 s pulse of 21% $O_2$ (green), followed by a 40 s pulse of 7% $O_2$ (black), followed by a 40 s pulse of 21% $O_2$ (green). For violin plots, dots represent individual worms, solid lines show medians, and dotted lines show quartiles. *$p$<0.05, **$p$<0.01, ***$p$<0.001, ****$p$<0.0001, Friedman test with Dunn's post-test; only significant, adjacent comparisons are displayed. n = 18-21 worms per life stage. Data in A-B are from Fig. S4.

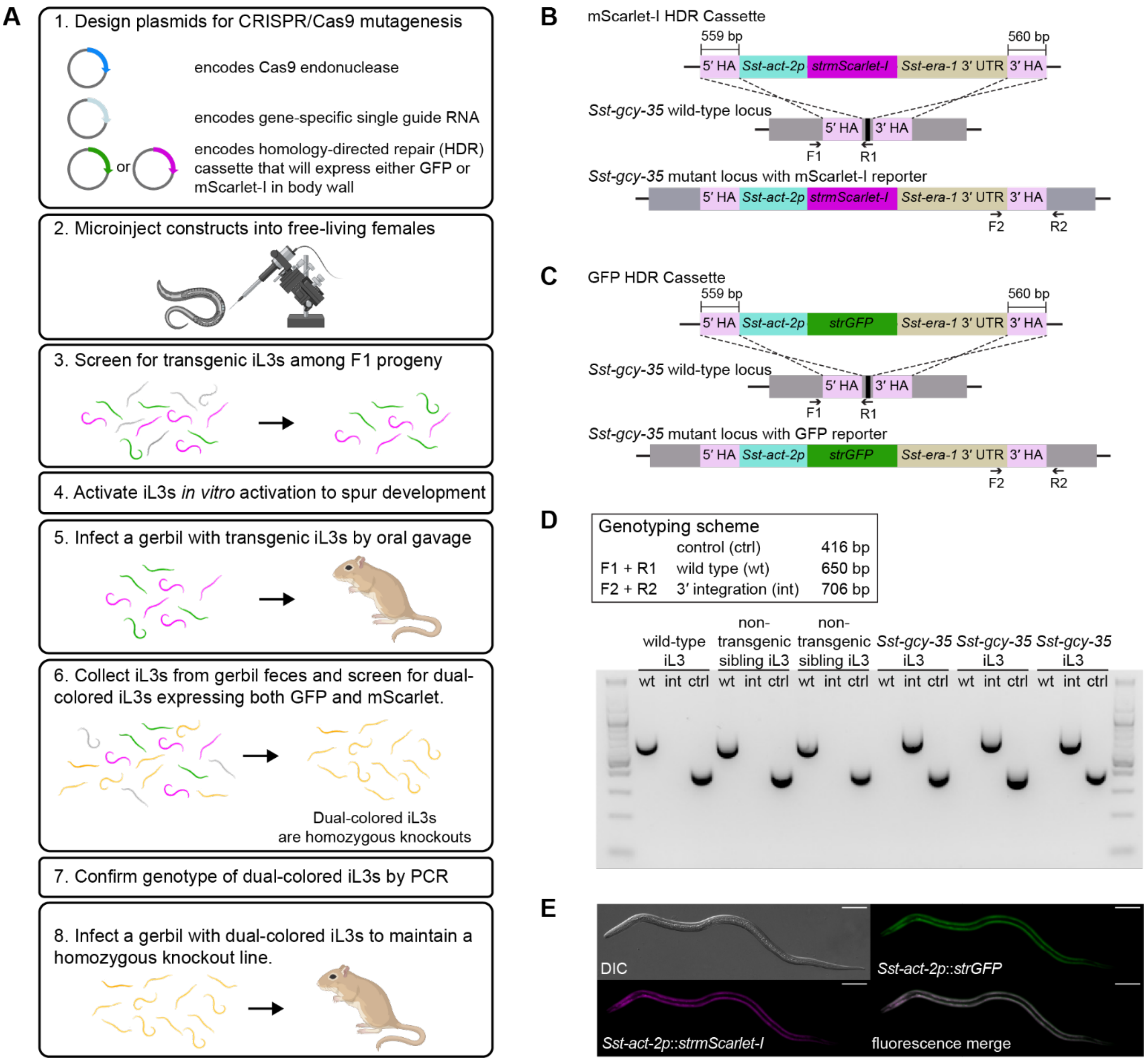


**Fig. S6. Strategy for generating a stable knockout line of *Sst-gcy-35* in *S. stercoralis*. A.** Approach for creating a *Sst-gcy-35* knockout line[1,12-14]. 1) The CRISPR components include plasmids encoding Cas9, the single guide RNA (sgRNA), and the homology-directed repair (HDR) cassette. The HDR cassette has a fluorescent reporter gene driven by a body-wall muscle promoter situated between 5′ and 3′ homology arms (HAs). 2) Plasmids are microinjected into the gonads of FLFs; half are injected with an HDR cassette that encodes GFP (green) and half are injected with an HDR cassette that encodes mScarlet-I (magenta). 3) After 5-7 days, the iL3 progeny of the injected FLFs are screened and worms with GFP or mScarlet-I expression throughout their body wall are isolated. 4) Transgenic iL3s are activated *in vitro* and are 5) introduced to a gerbil via oral gavage. 6) Worms are collected from gerbil feces; following sexual reproduction between FLFs and FLMs, a subset of the resultant iL3s will express both GFP and mScarlet-I (gold). 7) Dual-color iL3s are expected to be homozygous knockouts, which is confirmed by genotyping. 8) Homozygous knockouts can be selectively introduced into a gerbil, thus generating a stable line. **B-C.** HDR cassettes, wild-type locus, and mutated locus for *Sst-gcy-35*. The 5′ HA (559 bp, pink) and the 3′ HA (560 bp, pink) flank the CRISPR target site on the wild-type locus. The HDR cassette contains the *Sst-act-2* promoter, either *Strongyloides*-codon-optimized (*str*) *mScarlet-I* (**B**) or *GFP* (**C**), and the *Sst-era-1* 3′ UTR. Following integration, the HDR cassette will be inserted into the *Str-gcy-35* gene. Primers F1 and R1 amplify the wild-type locus; primers F2 and R2 amplify the mutated *Sst-gcy-35* locus. Black line indicates CRISPR target site. **D.** Representative agarose gel for single-worm genotyping. Six worms were genotyped: one wild-type iL3, two non-transgenic sibling iL3s (*i.e.,* worms isolated from the first infected gerbil that displayed no fluorescence), and three dual-color *Sst-gcy-35* iL3s. Primers F1 and R1 amplify a 650 bp product from the wild-type (wt) locus. Primers F2 and R2 amplify a 706 bp product from the mutant locus containing an integration event (int). A control PCR reaction (416 bp) confirms the presence of genomic DNA (ctrl). **E.** Dual expression of GFP and mScarlet-I in the body wall of an *Sst-gcy-35* iL3. Panels show differential interference contrast (DIC; top left), expression of the *Sst-act-2p*::*strGFP* reporter in green (top right), expression of the *Sst-act-2p*::*strmScarlet-I* reporter in magenta (bottom left), and the fluorescent merge of both reporters (bottom right). Head is to the left; scale bar is 50 µm.

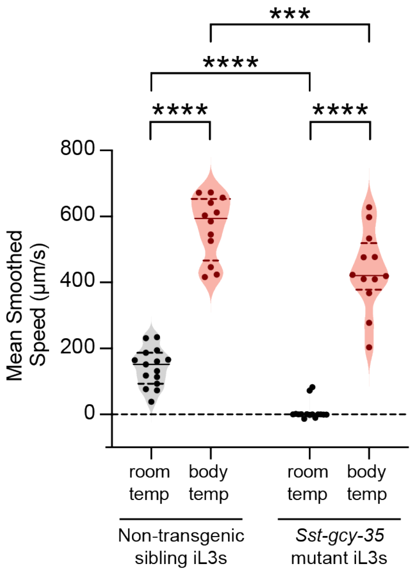


**Fig. S7. *Sst-gcy-35* iL3s are capable of rapid movement with a heat stimulus.** *S. stercoralis* control and *Sst-gcy-35* iL3s were introduced to agar plates either at room temperature (~21°C, black) or host body temperature (~34-37°C[15], red). Graph shows the mean smoothed speed over a time course of 30 s. Dots represent individual worms, with medians shown by solid lines and quartiles shown by dotted lines. Negative values indicate reverse movement. ****p*<0.001, *****p*<0.0001, ordinary two-way ANOVA with uncorrected Fisher's LSD post-test. n = 12-15 worms per genotype and condition.

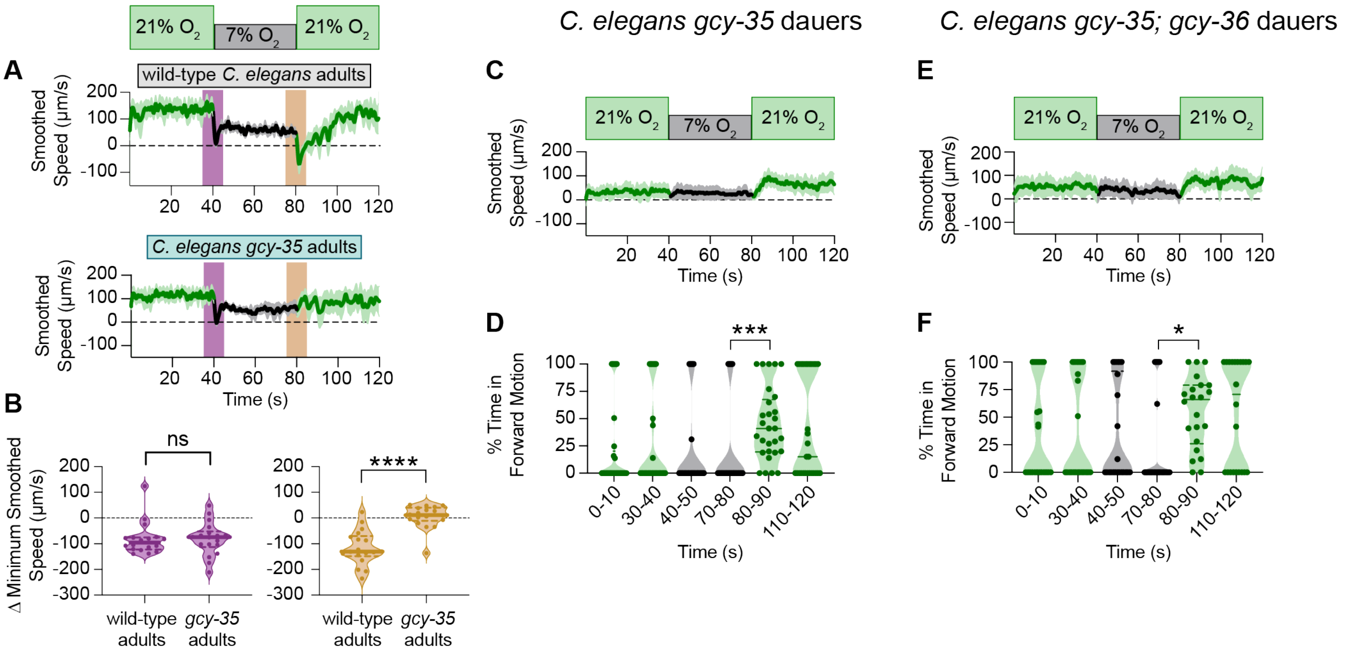


**Fig. S8. sGC repertoires sculpt $O_2$ responses. A-B.** *C. elegans gcy-35* adults fail to mount $O_2$-upshift-associated behaviors seen in wild-type *C. elegans* adults. **A-B.** Response of wild-type vs. *Cel-gcy-35* adults to acute $O_2$ shifts. **A.** Graphs of smoothed speed. Bold lines show mean smoothed speed; shading represents the 95% confidence interval. **B**. Violin plots show the difference in minimum speed in the 5 s before and after the downshift (purple) or upshift (yellow). For A, negative values indicate reverse movement. **C-D.** *C. elegans gcy-35* dauers respond to $O_2$ upshifts. Graph in C is as described in A. Graph in D shows the percentage of time spent in forward motion during the first and last 10 s of each gas pulse. **E-F.** *C. elegans gcy-35; gcy-36* dauers respond to $O_2$ upshifts. Graphs are as described in C-D. For A-F, worms were exposed to a 40 s pulse of 21% $O_2$ (green), followed by a 40 s pulse of 7% $O_2$ (black), followed by a 40 s pulse of 21% $O_2$ (green). For violin plots, dots represent individual worms, solid lines show medians, and dotted lines show quartiles. For B, ****$p<0.0001$, Mann-Whitney test. For D and F, *$p<0.05$, ***$p<0.001$, Friedman test with Dunn's post-test; only significant, adjacent comparisons are displayed. n = 18-29 worms per genotype and life stage. Data in A are from Fig. S5; data in C-D are from Fig. 4.

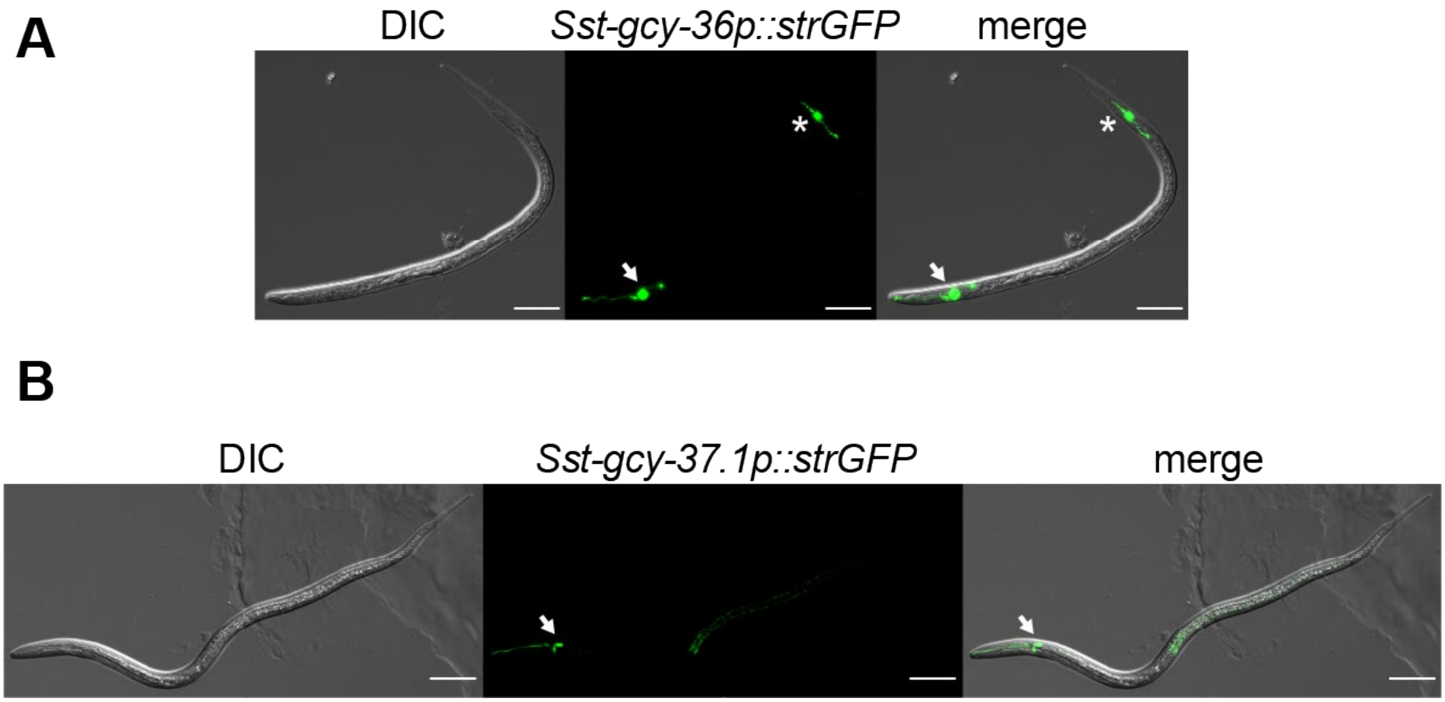


**Fig. S9. Transcriptional reporters for the *S. stercoralis* sGCs reveal neuronal expression patterns.** *Sst-gcy-36* (**A**) and *Sst-gcy-37.1* (**B**) are expressed in a subset of head and tail neurons that are anatomically and positionally similar to the $O_2$-sensing neurons in *C. elegans*. Left, differential interference contrast (DIC); center, GFP expression driven by the *Sst-gcy-36* promoter (**A**) or the *Sst-gcy-37.1* promoter (**B**); right, DIC and fluorescence merge. Head is to the left; scale bar is 50 µm.

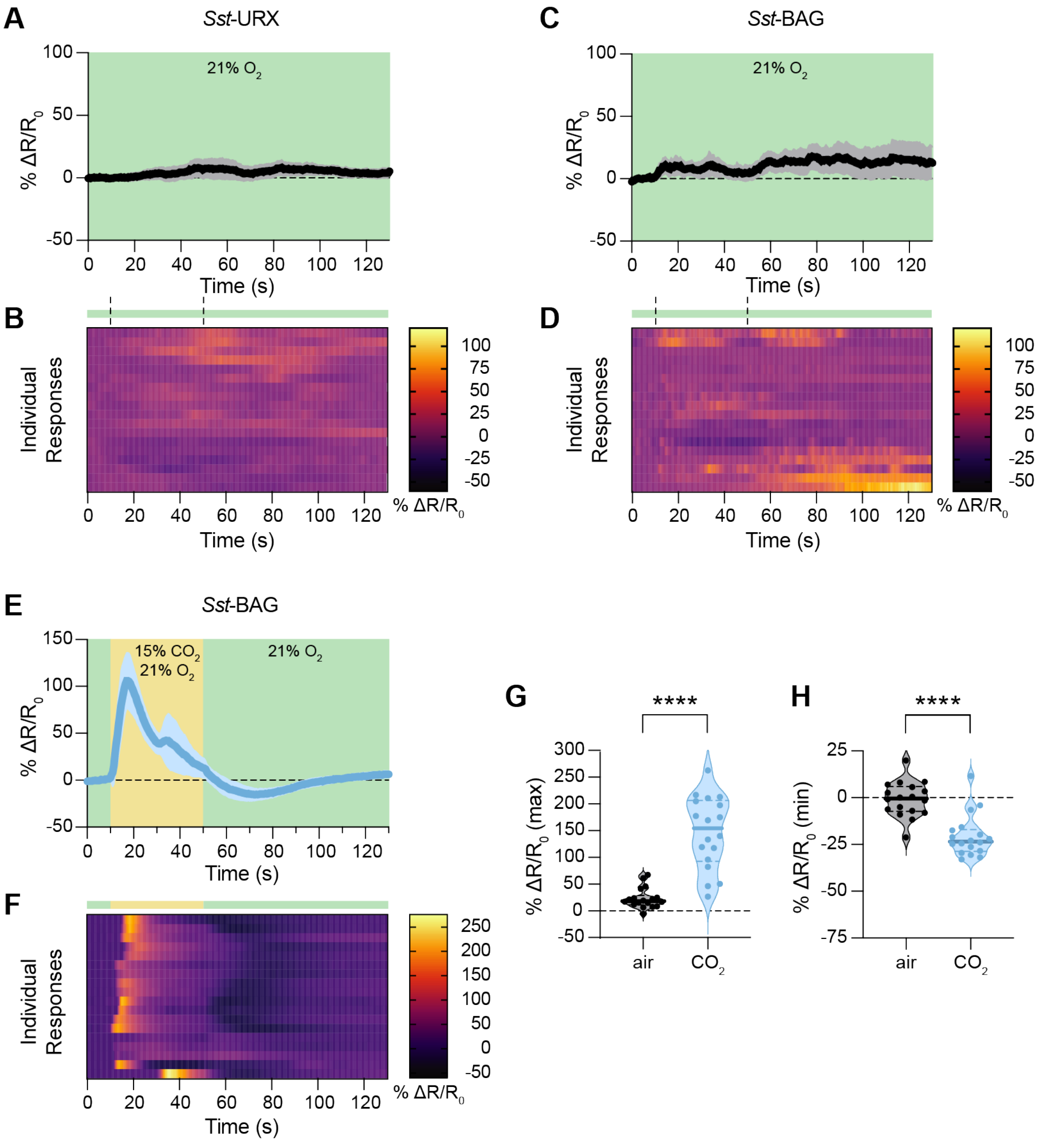


**Fig. S10. Calcium imaging controls in the *S. stercoralis Sst*-URX and *Sst*-BAG neurons. A-D.** *Sst*-URX and *Sst*-BAG neurons do not respond to successive pulses of 21% $O_2$. iL3s expressing GCaMP7f and mScarlet in either *Sst*-URX (**A-B**) or *Sst*-BAG (**C-D**) were exposed to a 40 s pulse of 21% $O_2$ (green), followed by a 40 s pulse of 21% $O_2$ (green), followed by an 80 s pulse of 21% $O_2$ (green); only the last 10 s of the initial pulse are displayed (t = 0-10 s). Graphs show the change in fluorescence intensity relative to baseline (% $\Delta R/R_0$); bold lines show means, and shading represents 95% confidence intervals (**A, C**). Heatmaps show the neuronal responses of individual worms (rows) throughout the assay duration (**B, D**). Bar indicates the stimulus paradigm; dashed lines indicate switches between $O_2$ tanks. n = 18 worms per genotype. **E-H.** *Sst*-BAG neurons are activated by $CO_2$. iL3s expressing GCaMP7f and mScarlet in *Sst*-BAG were exposed to a 40 s pulse of 0% $CO_2$ and 21% $O_2$ (green), followed by a 40 s pulse of 15% $CO_2$ and 21% $O_2$ (yellow), followed by an 80 s pulse of 0% $CO_2$ and 21% $O_2$ (green); only the last 10 s of the initial pulse are displayed (t = 0-10 s). Graphs show the change in fluorescence intensity relative to baseline (% $\Delta R/R_0$); bold line shows mean response, and shading represents the 95% confidence interval (**E**). Heatmap shows the neuronal responses of individual worms (rows) throughout the assay duration (**F**). Bar indicates the stimulus paradigm, with the $CO_2$ pulse in yellow. **G.** The maximum neuronal responses from t = 10-50 s (blue) are increased in worms exposed to 15% $CO_2$ and 21% $O_2$, when compared to maximum responses from worms exposed to 0% $CO_2$ and 21% $O_2$ (black). **H.** The minimum neuronal responses from t = 50-90 s (blue) are decreased in worms exposed to 15% $CO_2$ and 21% $O_2$, when compared to minimum responses from worms exposed to 0% $CO_2$ and 21% $O_2$ (black). ****$p$<0.0001, Welch's t test (**G**) or Mann-Whitney test (**H**). n = 18 worms.

### SUPPLEMENTAL TABLE

**Table S1. Oligonucleotides used in this study.**

| Oligonucleotide set (5' to 3') | Product size (base pairs) | Notes |
|---|---|---|
| **SG78 (F)**[12]**:** GTATTCCCTTCTATTGTTGGAAGACC **SG80 (R)**[12]**:** CCTTCATAGATTGGTACAGTGTGAG | 416 | Amplifies *Sst-act-2* exon 1; gDNA control ("ctrl") (Fig. S6B-D). |
| **BW40 (F)**: ACTTTTAATGAAAAATTGATACCATCTTTAGG **BW41 (R)**: CCATAAGCTTCCCAAATCTCTTCTATTGG | 650 | Amplifies wild-type ("wt") *Sst-gcy-35* locus (F1+R1, Fig. S6B-D). |
| **RP30 (F)**[16]: CGGTATATTTTACTTCAATGTGG **BW44 (R):** TGTTAATGGAAGTTCAGGATGC | 706 | Amplifies 3′ integration ("int") site in mutated *Sst-gcy-35* locus (F2+R2, Fig. S6B-D). |
